# CRISPR screens for lipid regulators reveal a role for ER-bound SNX13 in lysosomal cholesterol export

**DOI:** 10.1101/2021.05.10.443492

**Authors:** Albert Lu, Frank Hsieh, Carlos Enrich, Suzanne R. Pfeffer

## Abstract

We report here two genome-wide CRISPR screens carried out to identify genes that when knocked out, alter levels of lysosomal cholesterol or bis(monoacylglycero)phosphate. In addition, these screens were also carried out under conditions of NPC1 inhibition to identify modifiers of NPC1 function in lysosomal cholesterol export. The screens confirm tight co-regulation of cholesterol and bis(monoacylglycero)phosphate levels in cells, and reveal an unexpected role for the ER-localized, SNX13 protein as a negative regulator of lysosomal cholesterol export. In the absence of NPC1 function, SNX13 knockout decreases lysosomal cholesterol, and is accompanied by triacylglycerol-rich lipid droplet accumulation and increased lysosomal bis(monoacylglycero)phosphate. These experiments provide unexpected insight into the regulation of lysosomal lipids and modification of these processes by novel gene products.

**SUMMARY:** Genome-wide CRISPR screens carried out with and without NPC1 function identify shared pathways that coordinately control lysosomal cholesterol and bis(monoacylglycero)phosphate. ER-localized SNX13 protein plays an unexpected regulatory role in modifying NPC1 function to regulate cellular cholesterol localization.

## INTRODUCTION

Cellular lipid homeostasis is maintained by complex and dynamic, inter-organelle communication processes that coordinate uptake, biosynthesis and degradation of over 1000 lipid species. Among all the organelles involved in lipid regulation, the lysosome plays a central role. Lysosomes are the final station at which endocytosed lipoprotein particles and membranes derived from intra-lumenal budding and autophagy undergo a series of degradative reactions to yield unesterified cholesterol and other lipid precursors (Gruenberg, 2019; Ballabio and Bonifacino, 2020). Free cholesterol is then exported out of the lysosome and either recycled for *de novo* synthesis of biological membranes and other sterol products or esterified and stored in lipid droplets.

Loss-of-function mutations in gene products that mediate lipid turnover or lysosomal export lead to lethal cell toxicity, as seen in a number of inherited human diseases collectively known as lysosomal storage diseases (LSDs) (Ballabio and Bonifacino, 2020). One of the most studied LSDs is Niemann Pick type C (NPC) disease, caused by genetic defects in the lysosomal cholesterol transport system, Niemann Pick C1 and C2 proteins (NPC1 and NPC2). Cells from NPC patients accumulate cholesterol and glycosphingolipids in lysosomes, eventually leading to incurable neurodegeneration and premature death (Pentchev, 2004). Despite recent advances in the understanding of how transmembrane NPC1 and lumenal NPC2 coordinate their functions to export cholesterol from the interior to the outer-limiting membrane of the lysosome (Pfeffer, 2019), the precise molecular events that regulate this process and function downstream of NPC1 are still unclear (Das et al., 2014; Infante and Radhakrishnan, 2017).

An important player in the metabolic regulation of cholesterol is Bis(monoacylglycero)phosphate (BMP, also known as LBPA), a negatively charged glycerophospholipid with unique physicochemical properties that is found almost exclusively in intralumenal vesicles of multivesicular endosomes (MVEs; Gruenberg, 2020). Elevation of BMP levels occurs in many LSDs, including NPC, and BMP plays important roles in lipid catabolism and cholesterol egress from lysosomes (Chevallier et al., 2008). Indeed, Storch and colleagues have shown BMP activation of NPC2-mediated retrieval and transfer of cholesterol to NPC1 protein (reviewed in McCauliff et al., 2019). Moreover, in the absence of active NPC1, cells fed phosphatidylglycerol increase their BMP content which decreases their lysosomal cholesterol levels by a process that requires NPC2 (McCauliff et al., 2019). These data support the existence of NPC1-independent, relatively slow cholesterol export pathways. Nevertheless, it is still not known precisely how BMP is synthesized or degraded in cells.

Once cholesterol exits lysosomes, membrane contact sites between the lysosome surface and other juxtaposed compartments deliver cholesterol to the ER, a process that involves transit via the plasma membrane (Infante and Radhakrishnan, 2017). Because much remains to be learned regarding the mechanisms of cholesterol transport and its regulation, we performed genome-wide CRISPR screens to identify regulators of lysosomal cholesterol and BMP homeostasis. We repeated these screens under conditions of NPC1 inhibition to identify cellular components that may function in parallel with the NPC1 pathway to accomplish cholesterol export; such gene products might offer pathways to benefit patients with NPC disease. This strategy allowed us to confirm known components and pathways that regulate intracellular cholesterol transport and metabolism, and revealed many other previously unrecognized players. As one example, we show here that SNX13, a poorly characterized ER-resident inter-organelle tether, regulates lysosomal cholesterol export in an NPC1-independent manner. Remarkably, in the absence of NPC1 function that normally leads to massive lysosomal cholesterol accumulation, SNX13-depleted cells do not accumulate cholesterol in the lysosome and instead redistribute it to the plasma membrane and other compartments.

## RESULTS

### Genome-wide screens to identify regulators of endolysosomal cholesterol

We established two screening protocols to monitor changes in either cholesterol or BMP using fluorescently tagged Perfringolysin O* to label accessible cholesterol (Das et al., 2013) or a monoclonal antibody to detect BMP in K562 cells in conjunction with flow cytometry (Fig. 1A). Briefly, Cas9-expressing K562 cells were infected with libraries containing 10 sgRNA guides per gene and grown for 10 days before analysis for their content of cholesterol or BMP. In addition, both screens were also carried out in parallel under conditions in which the lysosomal cholesterol exporter, NPC1, was inhibited by addition of U18666A, to identify modifiers of NPC1 protein function (Lu et al., 2015). Cells were labeled and sorted by flow cytometry and cells displaying the highest or lowest 10% signals for each marker were collected and sequenced (Fig. 1A).

**Figure 1.**
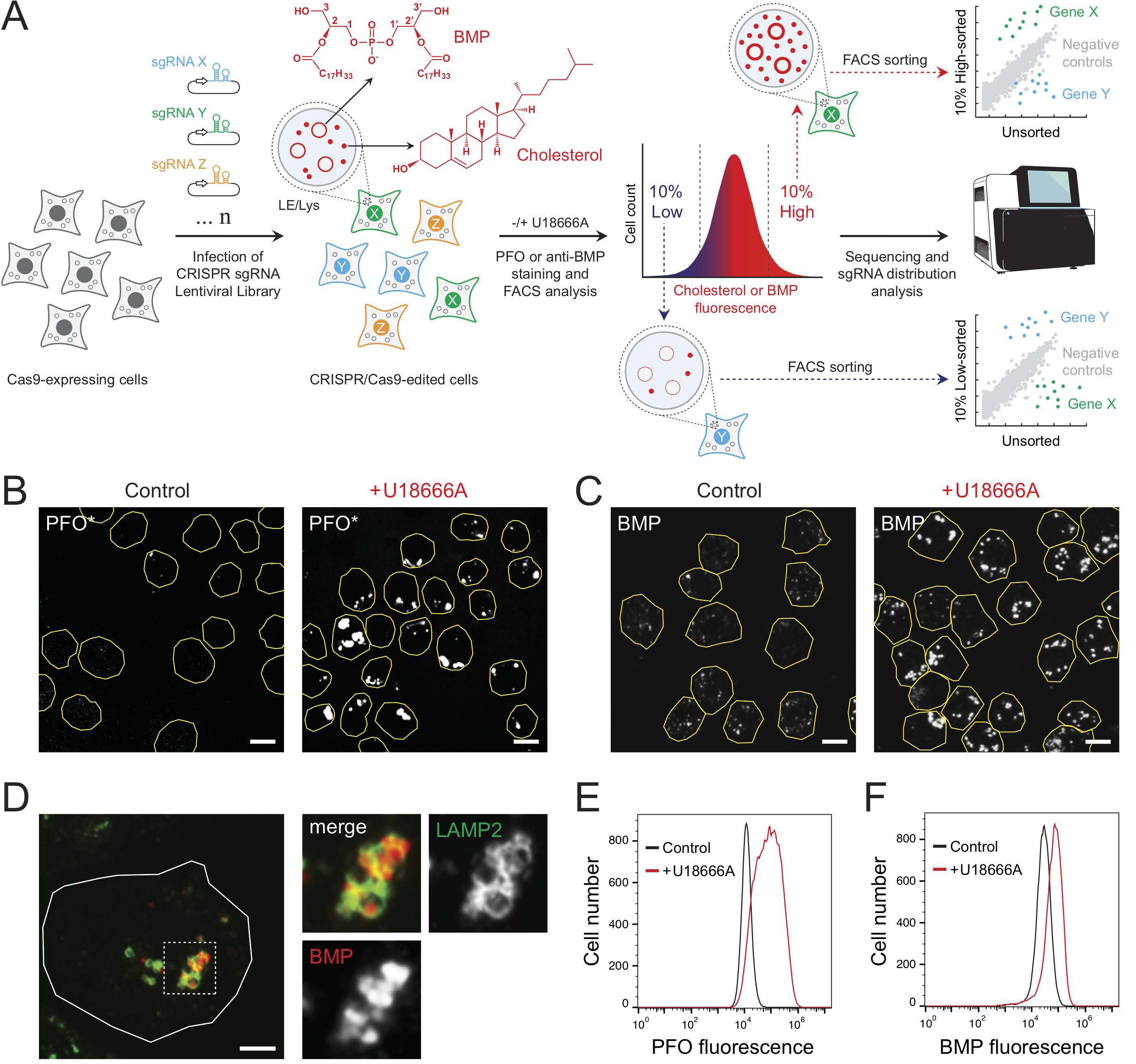
Flow cytometry based screen to identify cholesterol and BMP homeostatic regulators. (A) A Genome-wide CRISPR sgRNA lentiviral library (10 sgRNAs/gene) was used to infect Cas9-expressing human K562 cells such that after infection and selection, each cell expresses only a single sgRNA. CRISPR/Cas9-edited cells were then stained with either anti- BMP antibodies or fluorescently-conjugated, cholesterol binding Perfringolysin O* (PFO). Cholesterol and BMP (shown in red) accumulate in intra-lumenal vesicles (represented in red circles) of multivesicular endosomes and lysosomes (LE/Lys) upon treatment with the NPC1 inhibitor, U18666A. Cholesterol/BMP-stained cells were then sorted by flow cytometry and those showing the lowest 10% or highest 10% fluorescence (as indicated) were collected and compared with unsorted cells. Finally, sgRNAs were quantified by deep sequencing of sorted populations and compared to sgRNA counts in unsorted cells. (B) Immunofluorescence of K562 cells labeled with fluorescent PFO* ± U18666A for 16 hr. Cell boundaries (yellow) are shown and were determined by mCherry expression due to lentiviral transduction. (C) K562 cells stained with anti-BMP antibodies and circled as in (B). (D) K562 cells with anti-LAMP2 (green) and anti-BMP antibodies (red) as indicated. (E and F) Flow cytometry of cells labeled with either fluorescent PFO* or anti-BMP as in panels B and C, ±U18666A.

As shown in Fig. 1B, wild type K562 cells showed little PFO* staining unless NPC1 was inhibited using U18666A. Moreover, endolysosomal cholesterol accumulation seen with U18666A was accompanied by increased BMP that was localized (as expected) in LAMP2- positive structures (Fig. 1C,D). These phenotypes were easily scored by flow cytometry (Fig. 1E,F) where U18666A treated cells were easily resolved by their cholesterol or BMP content.

Figure 2 presents the compiled results monitoring the effects of gene depletions on cholesterol, BMP, and both of those lipids with and without U18666A, all carried out twice. Additional comparisons can be found in Supplemental Figure 1; gene hit subcellular localizations and associated metabolic pathways are presented in Supplemental Figures 2 and 3. Genes in the upper left and right quadrants of Fig. 2 increase cholesterol upon knockout; genes in the right top and bottom quadrants increase BMP upon knockout. Importantly, as expected, the strongest hits triggering cholesterol accumulation under control conditions were NPC1 and MYLIP, a ubiquitin ligase that regulates LDL receptor levels and would be expected to increase cholesterol uptake upon deletion. Similarly, knockdown of LDLR or the LDLRAP1 LDLR-specific endocytic adaptor decreased endolysosomal cholesterol and BMP, both with and without U18666A treatment. This recapitulates prior studies showing that cholesterol accumulation in lysosomes is accompanied by increases in antibody-detected BMP (Gruenberg, 2020). Knockdown of the IGF2R mannose 6-phosphate receptor had a much greater effect on BMP levels than cholesterol levels. This suggests that cells may upregulate BMP synthesis to compensate for a defect in lysosomal enzyme delivery; alternatively, it may reflect the requirement for a mannose 6-phosphorylated enzyme(s) to carry out BMP degradation. It is also of interest that most of the hits were located in the upper left or lower right quadrants of the graphs, with no hits increasing cholesterol while simultaneously decreasing BMP and conversely, no hits increasing BMP and decreasing cholesterol.

**Figure 2.**
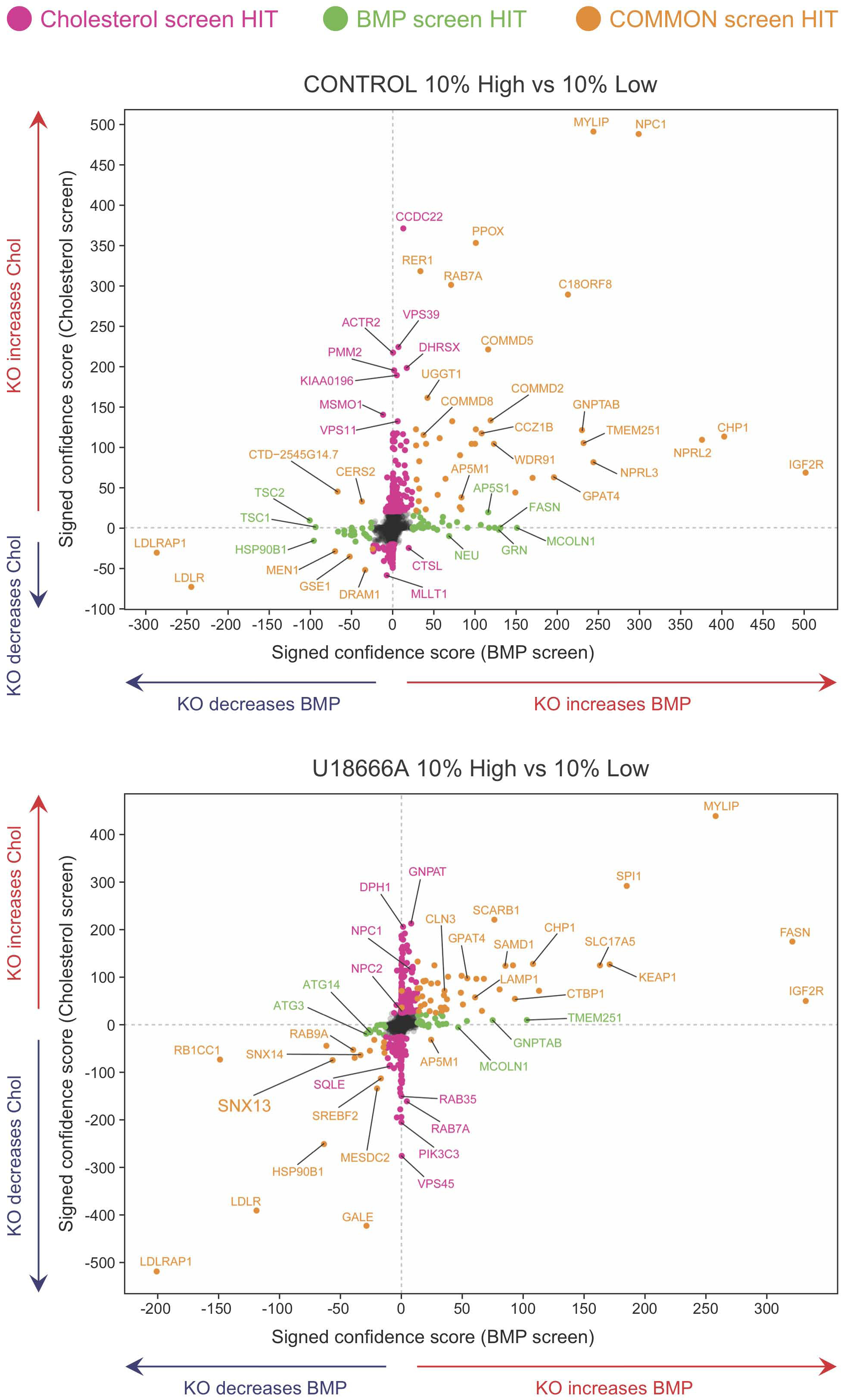
Bivariate analyses comparing hits in relation to changes in cholesterol and BMP. Indicated genes are presented in relation to their phenotypes determined by PFO* and BMP detection and flow cytometry, presented as signed confidence scores. Genes that appeared in both analyses are shown in goldenrod; those seen only in the cholesterol or BMP screens are shown in pink or green, respectively. Note the absence of hits in the upper left and lower right quadrants. The panels represent hits discovered in the top 10% versus bottom 10% comparisons, ± U18666A (upper versus lower graphs) as indicated.

**Figure 3.**
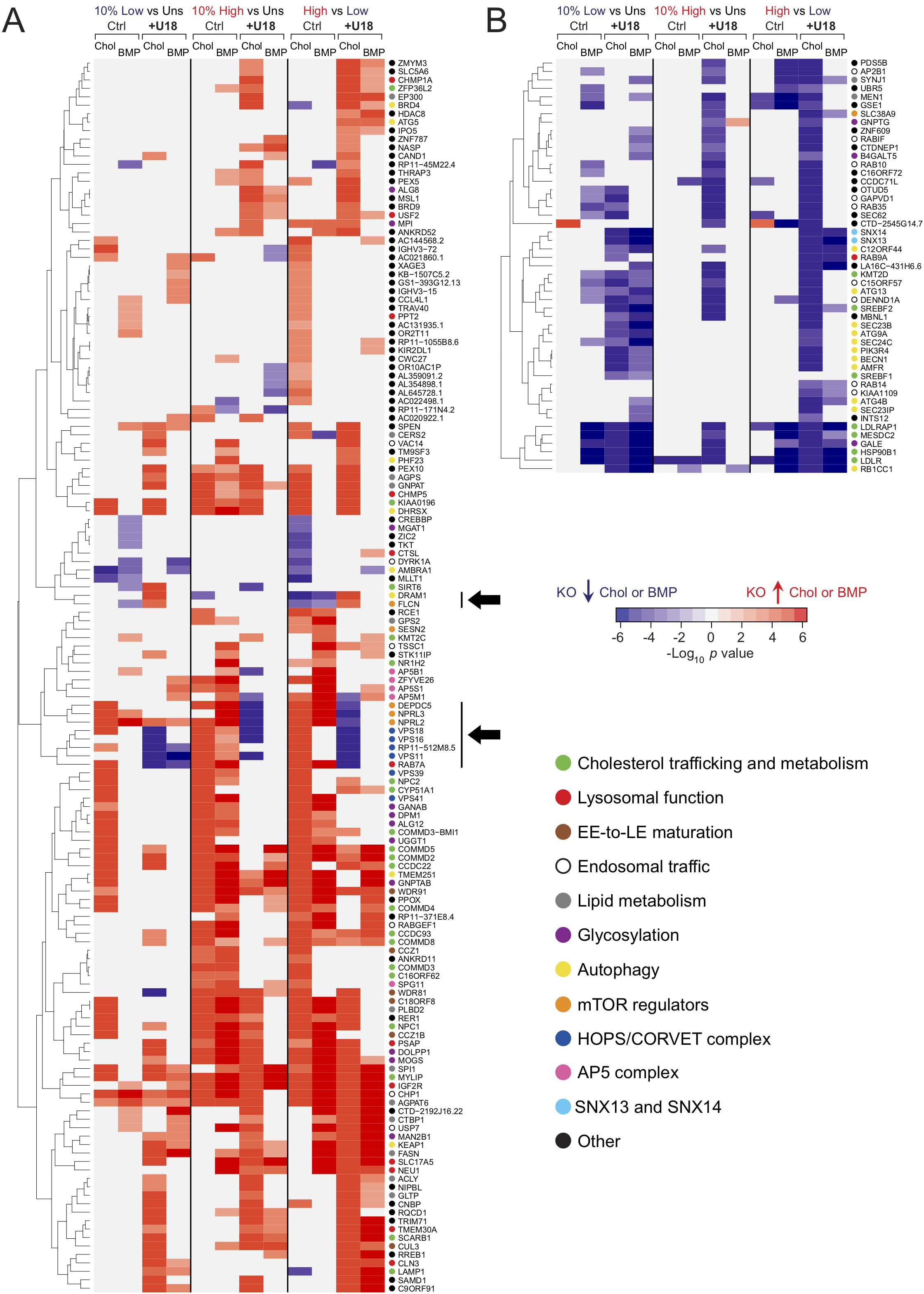
Hierarchical clustering analysis comparing 10% low and 10% high populations. Two main categories of genes, those increasing (A) or decreasing (B) cholesterol (Chol) and/or BMP are shown. For each category, the enrichment for cholesterol or BMP is shown as a heat map value. Responses under control (Ctrl) and U18666A (U18) conditions are also shown. The three types of screen data analysis performed (10% High vs Unsorted, 10% Low vs Unsorted and 10% High vs 10% Low) are indicated at the top. Only genes that were detected in both cholesterol and BMP screens are shown. Color intensity is presented using a log scale heat map proportional to the signed P value, as indicated. Individual hits in the high 10% pool are generally enriched in that population but may also be detected in the low 10% pool, explaining why red hits appear more populated on the 10% High vs Uns (unsorted) columns in (A) and blue hits on the 10% vs Unsorted columns in (B). In general the behavior is consistent. Black arrows indicate hits that upon deletion yielded opposite phenotypes in control versus U18666A conditions. Individual gene functions are shown using colored circles as indicated.

The presence of U18666A to inhibit NPC1 should increase lysosomal cholesterol accumulation yet knockout of SNX13 and to a lower extent, SNX14, decreased both BMP and cholesterol in the K562 cells used for the screen (Fig. 2, lower panel, lower left quadrant). Snx14 is an ER-resident protein that accumulates within a subdomain of the ER surrounding lipid droplets in oleic acid-fed cells (cf. Henne et al., 2015; Datta et al., 2019, 2020; Ugrankar et al., 2109). Further studies on the role of SNX13 in cholesterol regulation are presented below.

These screens generated an enormous amount of data and were analyzed by multiple means. Shown in Fig. 3 is a hierarchical classification of 195 hits that were identified in both cholesterol and BMP screens, displayed in terms of their phenotypes. Comparing genes enriched in the 10% high or 10% low groups, one would expect enrichment on one side or the other. Similarly, comparison of the behavior of each gene reveals hits that respond the same way when cholesterol or BMP phenotypes are scored. Two main categories of hits were revealed: hits that either increase (Fig. 3A) or decrease (Fig. 3B) cholesterol or BMP upon CRISPR deletion. A minor, third category of hits that yielded opposite results in each category was also observed (see genes marked by black arrows in Fig. 3A). This analysis revealed coordinate behavior of RAB7 and the HOPS complex VPS11, VPS16 and VPS18 subunits, that when knocked out in control conditions led to increased cholesterol and BMP, but in the presence of U18666A, strongly decreased cholesterol. It is possible that loss of endosome fusion capacity leads to accumulation of cholesterol in much smaller structures. Alternatively, loss of these proteins may trigger multivesicular endosome exocytosis. Similarly, the entire GATOR1 complex, comprised of DEPDC5, NPRL2 and NPRL3 encoded subunits, negatively regulates mTOR signaling and also decreases cholesterol accumulation but only when NPC1 function is blocked. On the other hand, the positive mTOR regulator, folliculin (FLCN), which counteracts the activity of GATOR1, yields an opposite profile (see genes marked by black arrows in Fig. 3A). Zoncu and colleagues (Davis et al., 2021) have shown that NPC1 loss elevates mTORC1 signaling, and inhibition of mTORC1 can improve lysosome proteolytic capacity without correcting cholesterol accumulation. Here, loss of the negative regulatory activity of the GATOR1 complex may show a worse phenotype because of dysregulation of hyperactive mTORC1.

Autophagy-related genes dominated the group that decrease cholesterol and/or BMP (annotated with a yellow dot, Fig. 3B bottom right). Auto-phagocytosed membranes, derived from mitophagy, ER-phagy or lipophagy for example, represent a major source of lipid substrates in endo-lysosomes. Perhaps inhibition of autophagy decreases lysosomal catabolic burden and consequently, the amount of BMP needed for lipolytic reactions (Kolter and Sandhoff, 2005), as well as the levels of intra-lysosomal free cholesterol. Alternatively, under conditions of decreased autophagy, endolysosomes may be smaller and be more difficult to detect by flow cytometry analogous to knockout of HOPS subunits. Finally, cholesterol homeostasis regulators (green circles) including LDLR, LDLRAP1, and the LDLR chaperones HSP90B1 and MESDC2, as well as SREBF1 and 2 are also prominent in this category, consistent with their known roles in cholesterol homeostasis.

### Comparison of four genome-wide screens for cholesterol regulators

Several groups have recently reported genome-wide screens to identify cholesterol homeostatic regulators. Gruenberg and colleagues presented an siRNA screen that detected a connection with Wnt signaling (Scott et al., 2016). Trinh et al. (2020) measured surface LDLR levels; van den Boomen et al. (2020) employed a synthetic reporter to monitor SREBP transcriptional activation; and Chu et al. (2015) used amphotericin to kill cells that succeeded in plasma membrane cholesterol delivery. Use of different approaches will surely yield different overall hit profiles. Shown in Figure 4 is a comparison of three of these genome screens and the screens presented here. This comparison identified only two completely overlapping hits, LDLR and NPC1, highlighting the fact that different approaches and readouts impact hit discovery. Nevertheless, a notable number of genes discovered here were also detected by several of the previous screens when compared individually. For example, our screen analysis uncovered roles for PTDSS1, SNX13 and SNX14 (Fig. 4; see also Fig. S2 and S3), all of which were ranked among the top 104 hits identified in the screens performed by Trinh et al. (2020). Moreover, several factors that regulate LDLR trafficking, including different subunits of the AP2, Arp2/3 and CCC complexes or the ubiquitin ligase MYLIP, also appear in this comparison. Notably, only a relatively small number of genes are shared between previous screens but not ours (see gene names associated with a yellow dot in Fig. 4). Altogether, this comparative analysis confirms the comprehensive nature of the present screens and the importance of independent evaluations of lipid regulatory pathways.

**Figure 4.**
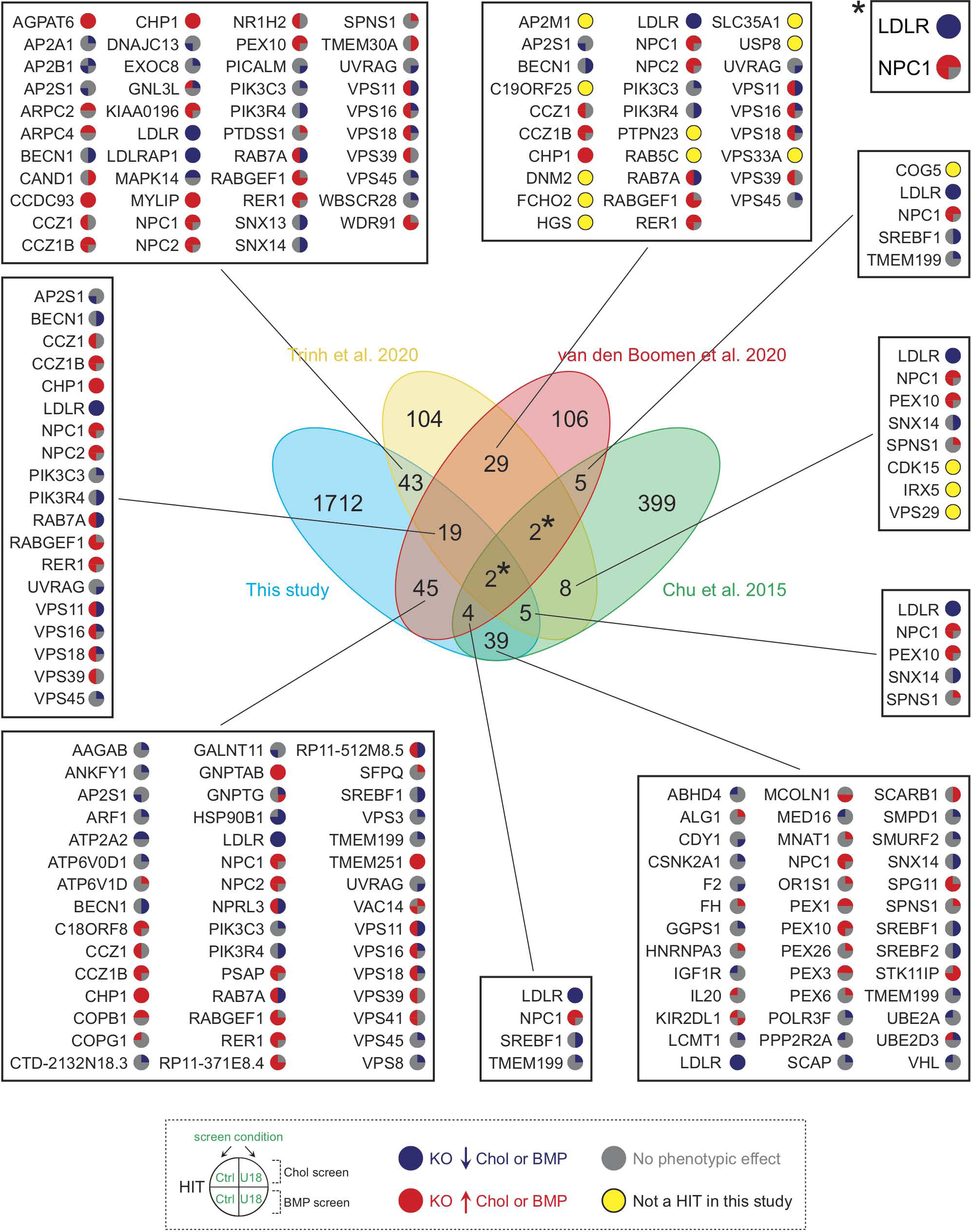
Comparison of hits from four independent studies analyzing cholesterol homeostasis. The central Venn diagram displays overlaps in hits from the indicated, referenced screens. Numbers in the non-intersecting areas indicate the number of hits identified in each study, whereas numbers in the intersecting areas correspond to overlapping hits between the different referenced screens. Specific gene hits from this screen are shown in adjacent boxes with colored circles according to an increase (red) or decrease (blue) in cholesterol or BMP. Genes that were identified here but only under one or another conditions are shown with a gray circle; those not detected at all are highlighted with a yellow circle.

To further illustrate the coverage of our screen data, we performed a functional interaction analysis of all hits identified in both cholesterol and BMP screens. This analysis revealed the most represented intracellular pathways and processes controlling cholesterol and BMP homeostasis (Fig. 5), with colors used to indicate decreases (blue) or increases (red) of these two lipids upon knockout, and with quadrants of the individual gene nodes showing results for each gene ± NPC1 function. As expected, a significant proportion of high-confidence interacting hits are involved in LDLR trafficking, SREBP pathway regulation or early/late endosomal function. Other well-represented functional gene clusters are those involved in autophagy and mTOR signaling, known to play critical roles in cellular lipid homeostasis (Thelen and Zoncu, 2017). A large number of interacting hits are involved in transcriptional regulation. Importantly, this analysis provides valuable information related to the coordinate function of individual subunits of protein complexes. We detected identical behavior for all subunits of the PRC2 chromatin remodeling complex and for the RISC complex (Fig. 5 bottom right and left, respectively). Similarly, all peroxisomal genes behaved similarly (Fig. 5 bottom left) although it is not yet clear what specific role they play in cholesterol and BMP regulation. Genes involved in LDLR recycling are all red (Fig. 5 upper left), suggesting, paradoxically, that cholesterol accumulates when LDLR cannot recycle. Perhaps this reflects trafficking of another protein(s) that facilitates cholesterol egress or increased receptor degradation followed by LDLR gene up- regulation. Unexpectedly, glycosylation genes are also mostly red (Fig. 5 top middle), possibly highlighting the importance of glycosylation for folding and assembly of membrane proteins that function in cholesterol export. Finally, while complexes involved in late endosomal maturation, such as the RAB7 GEF MON/CCZ1/C18ORF8 (van den Boomen, 2020) or the recently described WDR91/81 complex (Casanova and Winckler, 2017), show increased cholesterol or BMP accumulation upon knockout, endosome fusion complexes such as HOPS and Rab7 itself are mixed (Fig. 5 middle left; see Fig. S2): increasing cholesterol and BMP on knockout and decreasing cholesterol and BMP on knockout with U18666A. It is possible that this phenotypic difference may reflect the different sized endolysosomal compartments generated under the two conditions, and a possible up-regulation in MVE exocytosis in the presence of U18666A compound (Strauss et al., 2010).

**Figure 5.**
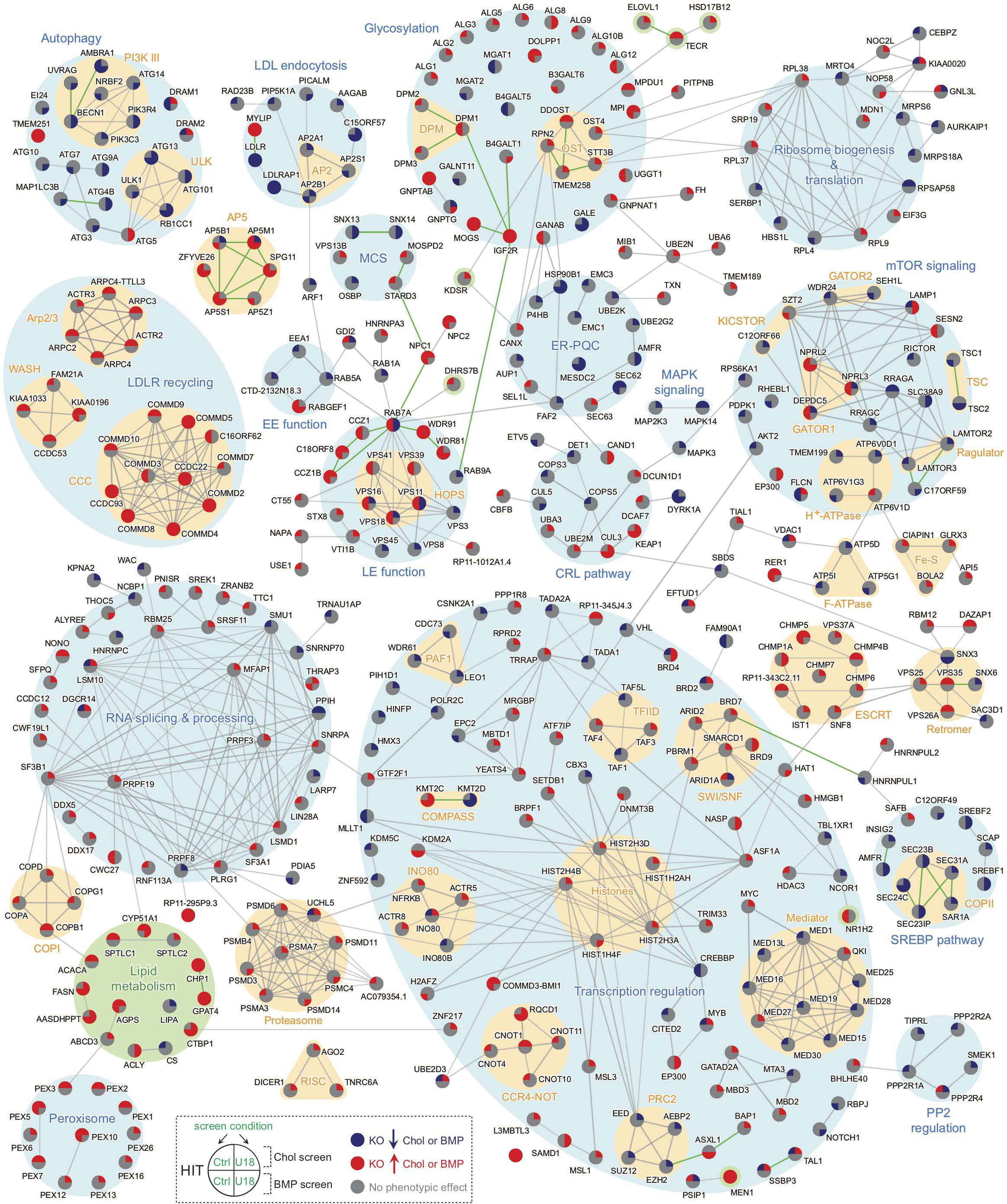
Functional network analysis of hits that interact according to STRING analysis. Genes are annotated using colored circles that show increases (red) or decreases (blue) in cholesterol (upper left quadrant), BMP (lower left quadrant), cholesterol +U18666A (upper right quadrant), or BMP + U18666A (lower right quadrant) as indicated. Clusters of functional categories are highlighted in light blue. Lipid metabolism-related hits are highlighted in green. Individual clusters highlighted in tan are known interacting protein complexes. Green edges are manually curated, reported interactions. MCS= Membrane contact sites, EE= Early endosome, LE= Late endosome, PP2= Protein phosphatase 2.

We also classified our hits according to specific lipid metabolic pathways (Fig. S3). In general, the majority of hits that catalyze the biosynthesis of multiple lipid species appear to increase either cholesterol or BMP or both, underscoring the complex interplay between these metabolic pathways. Our approach recovered key enzymes required for de novo cholesterol biosynthesis such as HMGCR, MSMO1, FDFT1, SQLE, and CYP51A1, and also identified the SREBP pathway almost in its entirety (Fig. S3 bottom right), including the recently described SREBP pathway regulator, C12ORF49 (Loregger et al., 2020; Bayraktar et al., 2020; Aregger et al., 2020). Moreover, identification of additional factors such as UBE2G2, which controls sterol- stimulated ubiquitylation and turnover of the rate-limiting enzyme HMGCR (Tan et al., 2019), or ZFP36L2 (Fig. S3 bottom right), a RNA-binding protein that promotes LDLR mRNA degradation (Adachi et al., 2014), further highlights the level of biological detail revealed by our screens. In most cases, their corresponding phenotypes were consistent with their respective biological functions in the context of cholesterol homeostatic regulation. Accordingly, while deletion of HMGCR resulted in elevated cellular cholesterol levels, UBE2G2 knockout cells showed the opposite phenotype; both genes appeared as hits only when NPC1 was pharmacologically blocked. Interestingly, a considerable number of transcription factors that control expression of lipid metabolism and lysosomal genes were also efficiently detected as hits in addition to SREBFs (see Figs. 5, S2 and S3), including liver X receptor member NR1H2 (Wang and Tontonoz, 2018), BRD4 (Sakamaki et al., 2017), SPI1 (Solomon et al., 2017), USF2 (Yamanaka et al., 2016) and the multi-subunit transcriptional co-activator, Mediator (Youn et al., 2016), once again validating the robustness of this approach.

### ER-localized SNX13 links to endolysosomes and lipid droplet domains

The goal of this screen was to identify modifiers of the NPC1 deletion phenotype, and SNX13 revealed itself as a gene that decreased PFO*-detected, accessible cholesterol and antibody detected-BMP in the presence of U18666A. Thus we carried out a cell biological characterization of the role of SNX13 in cholesterol regulation. Like SNX14, SNX13 is related to the single yeast MDM1 protein that mediates endoplasmic reticulum (ER) links to the yeast vacuole (Henne et al., 2015). SNX13 is a multi-domain containing protein comprised of PXA, RGS, PX and C-nexin domains (Fig. S4 F). In U2OS cells, we detected good co-localization of SNX13 with the ER localized VAP-A protein (Fig. S4 A). When NPC1 was inhibited, the protein remained in the ER and close apposition to lysosomes was detected (Fig. S4 B, RPE cell shown). Upon addition of oleic acid to induce lipid droplet formation, SNX13 redistributed to ER domains in contact with nascent lipid droplets (Fig S4 C), as shown recently for SNX14 (Datta et al., 2019 and Fig. S4 D). Detailed confocal microscopy (Fig. S4 E) showed intimate connections between SNX13 (green), BMP compartments (red) and lipid droplets (blue) under these conditions. SNX13 truncation analysis showed that the C-terminus is required for redistribution of SNX13 to lipid droplet forming domains (Fig. S4 F-J), analogous to SNX14 (Datta et al., 2019).

Immunofluorescence microscopy of U18666A-treated U2OS cells treated with SNX13 siRNA showed decreased cholesterol accumulation determined by PFO* staining compared with control siRNA or SNX14 siRNA-treated cells (Fig. 6 A-D). Specifically, SNX13-siRNA treated cells showed fewer PFO*-positive vesicles and vesicles were smaller in size (Fig. 6 E, F). Similar results were obtained using U18666A treated HeLa cells (Fig. 6 G-K).

**Figure 6.**
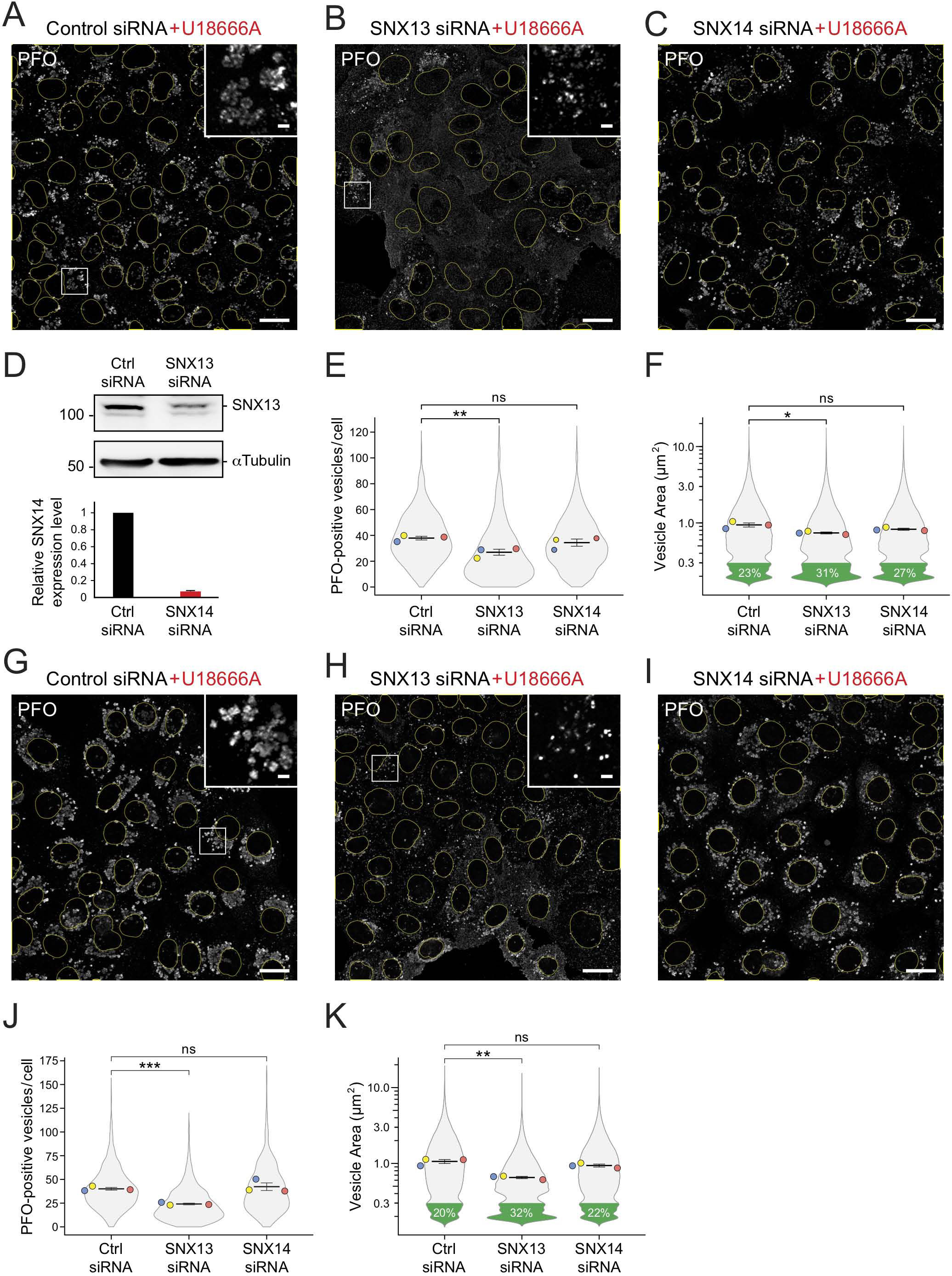
SNX13 depletion redistributes cellular cholesterol in the absence of NPC1 function. (A-C) Immunofluorescence microscopy of U2OS cells treated with U18666A for 16h and labeled with GST-PFO* detected using anti-GST primary antibodies, 72 hours after transfection with the indicated siRNAs. Scale bar, 10µm. Insets show enlargements of the boxed areas; scale bars, 2µm. (D) Immunoblot analysis (at the top) of SNX13 siRNA treated cells as in (A) and (B); 50µg cell extract was analyzed. Molecular weight markers are indicated at left in kD. Plot at the bottom shows relative SNX14 mRNA levels measured by qPCR using RNA isolated from control and SNX14 siRNA-treated U2OS cells from two independent experiments; error bar represents standard deviation. (E and F) Quantitation of PFO*-positive vesicles and area of vesicles determined by CellProfiler. Colored dots reflect means from independent experiments; >860 cells analyzed in each condition; significance was determined by unpaired *t* test; *, P < 0.05, **, P < 0.01. G-K, Analysis of HeLa cells as described for (A-F); >950 cells analyzed in each condition. Actual P values: (E: Ctrl siRNA vs SNX13 siRNA= 0.0079; Ctrl siRNA vs SNX14 siRNA= 0.16); (F: Ctrl siRNA+U18 vs SNX13 siRNA+U18= 0.016; Ctrl siRNA+U18 vs SNX14 siRNA+U18= 0.072); (J: Ctrl siRNA+U18 vs SNX13 siRNA+U18= 0.00035; Ctrl siRNA+U18 vs SNX14 siRNA+U18= 0.31); (K: Ctrl siRNA+U18 vs SNX13 siRNA+U18= 0.0019; Ctrl siRNA+U18 vs SNX14 siRNA+U18= 0.091).

In addition to decreasing lysosomal cholesterol, SNX13 depleted, U18666A-treated U2OS cells (Fig. 7A-D) and HeLa cells (Fig. S5 A-C) showed clear, NPC1-independent redistribution of cholesterol from endolysosomal compartments to the cell surface of non-permeabilized cells that was not seen in control siRNA and U18666A-treated cells (Fig. 7 A and Fig. S5 A) or SNX14 depleted cells (Fig. 6 C, I). This was again unexpected, as NPC1 is thought to be required upstream of cholesterol delivery to the cell surface. One possibility is that loss of SNX13 enhances formation of an endosomal-plasma membrane and/or ER-plasma membrane contact site that can bypass NPC1 (as seen by Höglinger et al., 2019). Importantly, depletion of SNX13 from NPC1 knockout HeLa cells yielded the same phenotype (Fig. S5 D), showing that this reflects a block in NPC1 function rather than an off target effect of U18666A.

**Figure 7.**
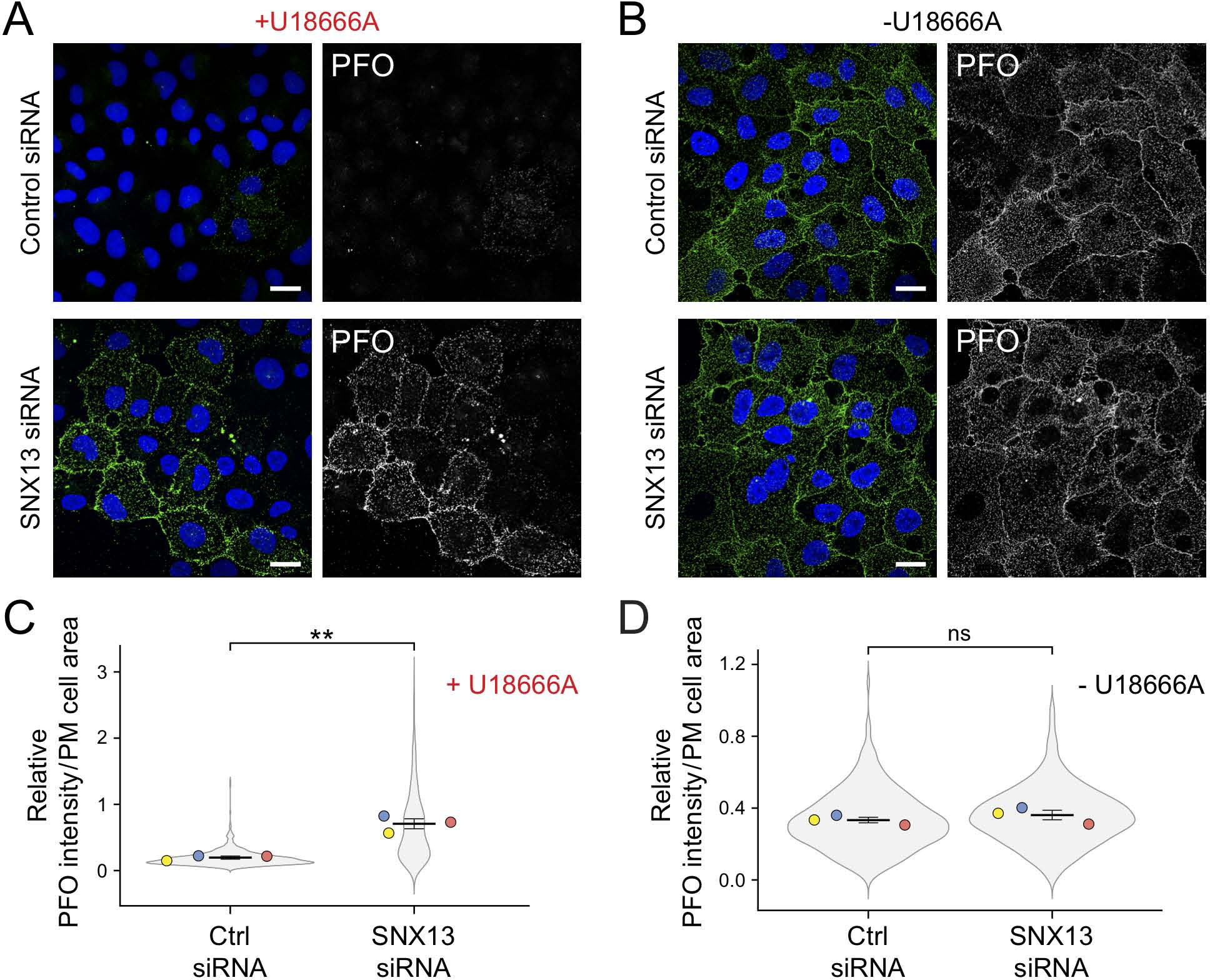
SNX13 depletion redistributes cholesterol to the cell surface in the absence of NPC1 function. (A and B) Immunofluorescence microscopy of U2OS cells treated with U18666A for 16 hours (A) or left untreated (B), and labeled in the absence of detergent permeabilization with GST-PFO* detected using anti-GST primary antibodies, 72 hours after transfection with the indicated siRNAs. (C and D) Quantitation of PFO* staining as a function of plasma membrane cell area in U18666A-treated (C) and untreated (D) cells, determined using CellProfiler. Colored dots reflect means from independent experiments; >630 cells analyzed in each condition; significance was determined by unpaired *t* test; **, P < 0.01. Scale bars, 20µm. Actual P values: (C: Ctrl siRNA+U18 vs SNX13 siRNA+U18= 0.0015); (D: Ctrl siRNA vs SNX13 siRNA= 0.209).

Zoncu and colleagues have shown that blocking NPC1 function hyperactivates mTORC1 and leads to accumulation of accessible cholesterol on the outer leaflet of lysosomes, highlighting a cholesterol transfer pathway from the ER to the lysosome outer leaflet (Lim et al., 2019; Davis et al., 2021). Somehow, loss of SNX13 enables cholesterol transfer to the plasma membrane despite the absence of NPC1 function. This phenotype suggests a role for SNX13 as a negative regulator of cholesterol transport from lysosomes to the cell surface, in coordination with NPC1.

SNX13 depletion, even in the absence of oleic acid supplementation, also increased the number of lipid droplets per cell, as monitored by LipidTOX staining (Fig. 8 A,B). This increase was seen whether or not cells were treated with the SANDOZ ACAT1 inhibitor, suggesting that these lipid droplets contain primarily triglycerides rather than cholesterol esters. Total lipids were analyzed by thin layer chromatography, which revealed elevated levels of both total triacylglycerol and free fatty acids, consistent with the increased, NPC1-independent, lipid droplet accumulation observed (Fig. 8 C,D).

**Figure 8.**
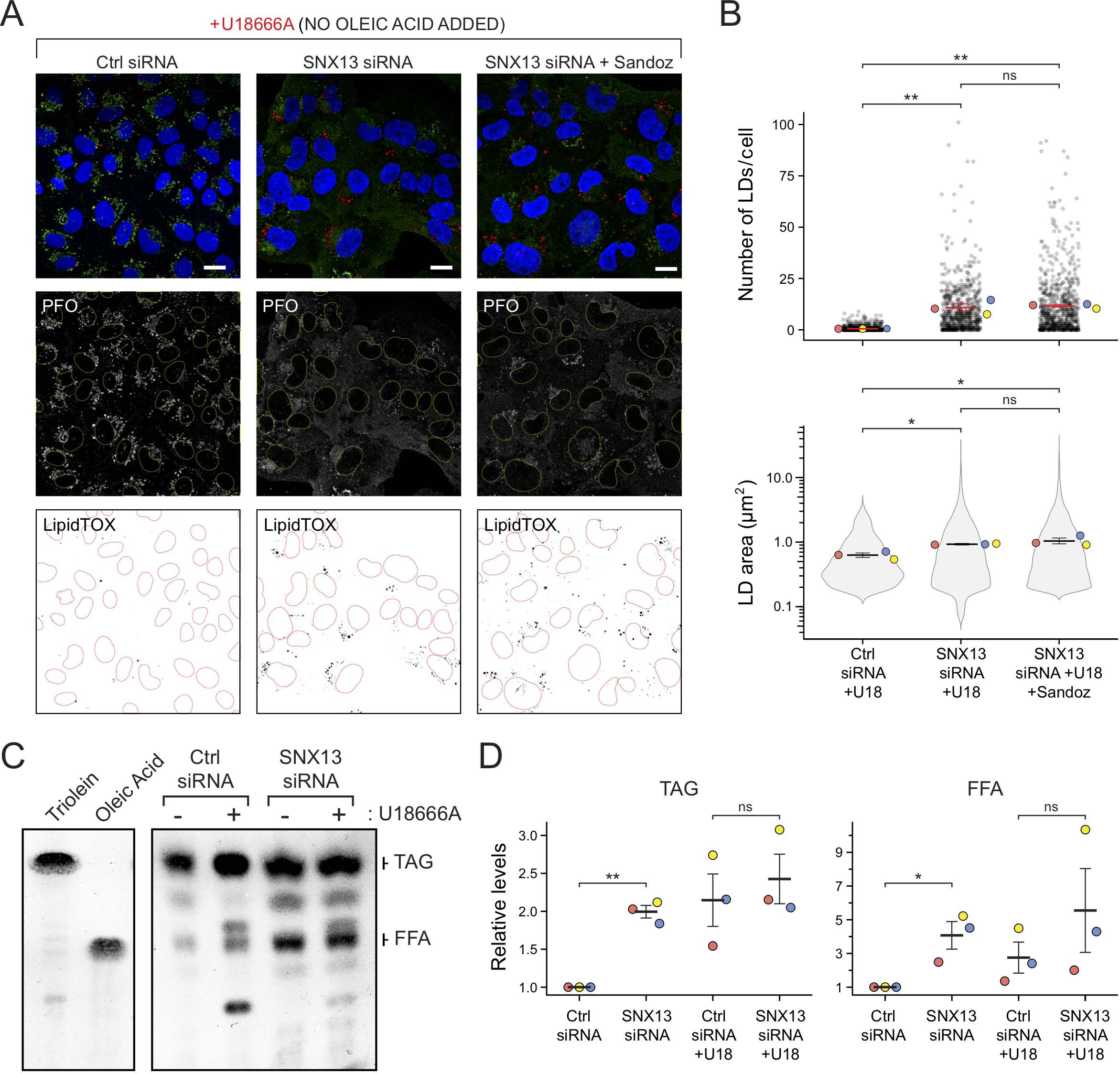
SNX13 depletion increases lipid droplet abundance in the absence of NPC1 function. (A) Immunofluorescence microscopy of U2OS cells treated with U18666A for 16h and labeled with GST-PFO* detected using anti-GST primary antibodies, 72 hours after transfection with the indicated siRNAs, without oleic acid addition. Cells were also stained with LipidTOX to highlight lipid droplets. One set of cells was also treated with the ACAT1 inhibitor Sandoz 53-035 as indicated. Scale bar, 10µm. (B) Number of lipid droplets (LDs) and their area determined by Cellprofiler for cells as in (A). Colored dots reflect means from independent experiments; >650 cells analyzed in each condition; significance was determined by multiple *t* test, Holm-Sidak method with α = 0.05. (C) Thin layer chromatography of lipid levels from cells treated as in (A). At left is shown the mobility of specific marker lipids. (D) Quantitation of triacylglycerol (TAG) and free fatty acid (FFA) levels determined by thin layer chromatography as indicated. P values were determined by multiple *t* test corrected by the Holm-Sidak method; *, P < 0.05, **, P < 0.01. Actual P values: (B, top: Ctrl siRNA+U18 vs SNX13 siRNA+U18= 0.0052; Ctrl siRNA+U18 vs SNX13 siRNA+U18+Sandoz= 0.0039; SNX13 siRNA vs SNX13 siRNA+Sandoz= 0.85); (B, bottom: Ctrl siRNA vs SNX13 siRNA= 0.0451; Ctrl siRNA vs SNX13 siRNA+Sandoz= 0.0144; SNX13 siRNA vs SNX13 siRNA+Sandoz= 0.382; (D, left: Ctrl siRNA vs SNX13 siRNA= 0.00683; Ctrl siRNA+U18 vs SNX13 siRNA+U18= 0.312); (D, right: Ctrl siRNA vs SNX13 siRNA= 0.0198; Ctrl siRNA+U18 vs SNX13 siRNA+U18= 0.351).

The initial flow cytometry screen indicated that SNX13 knockout had the strongest phenotype when NPC1 function was inhibited, and that knockout concomitantly decreased BMP staining (Fig. 2, Fig. S1). Yet immunofluorescence microscopy showed accumulation of antibody-detected BMP in NPC wild type, SNX13 siRNA-depleted cells (Fig. 9). This difference may be due to the fact that the screen relied on CRISPR deletion whereas our validation relies on siRNA; alternatively, it could reflect differences between K562 cells used in the screen and U2OS cells used in validation. Many studies have now shown that cells subject to CRISPR modification adapt in diverse ways (Cerikan et al., 2016; Diofano et al., 2020; Salanga and Salanga, 2021).

**Figure 9.**
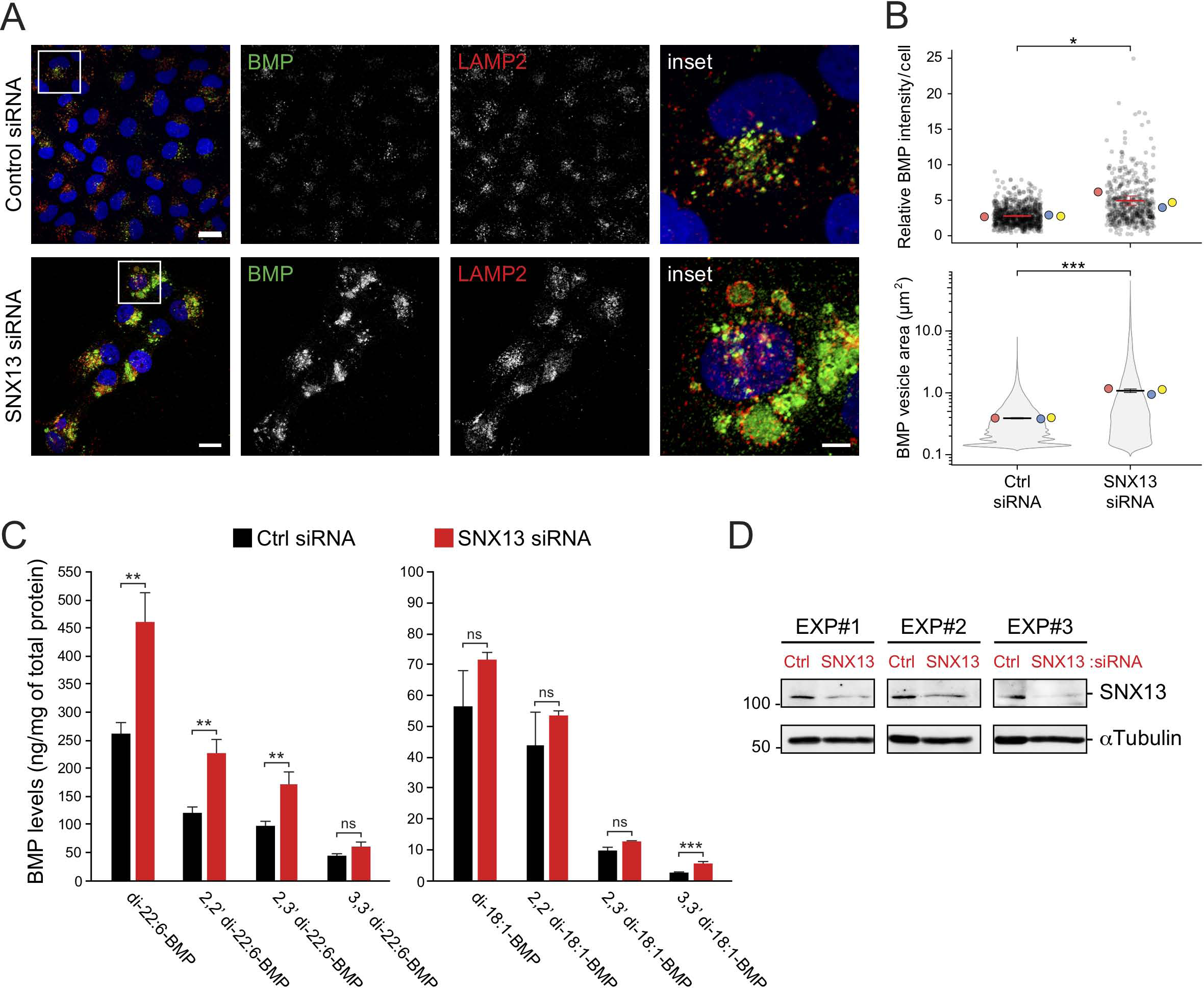
SNX13 depletion in NPC1 wild type, U2OS cells increases BMP levels. (A) Immunofluorescence microscopy of U2OS cells labeled with anti-BMP antibodies, 72h after transfection with the indicated siRNAs. Cells were also labeled with anti-LAMP2 antibodies as indicated. Scale bar, 20µm; in enlarged insets, 5 µm. (B) Quantitation of BMP-positive vesicle area and total BMP intensity per cell determined using Cellprofiler. Colored dots represent means from independent experiments; >480 cells analyzed in each condition; significance was determined by unpaired *t* test; *, P < 0.05,***, P < 0.001. (C) Mass spectrometric determination of BMP isoforms normalized to protein content from cells treated with the indicated siRNAs. Inset, western blot confirming SNX13 depletion from three independent experiments. Significance was determined by two-way ANOVA with Tukey’s post hoc test; *, P < 0.05, **, P < 0.01,***, P < 0.001. (D) Immunoblot analysis of SNX13 siRNA treated cells from the experiments in C; 50µg cell extract was analyzed. Molecular weight markers are indicated at left in kD. Actual P values: (B, top: Ctrl siRNA vs SNX13 siRNA= 0.0171); (B, bottom: Ctrl siRNA vs SNX13 siRNA= 0.000305); (C, left: di-22:6-BMP= 0.0024; 2,2’ di-22:6-BMP= 0.0018; 2,3’ di-22:6-BMP= 0.0025; 3,3’ di-22:6-BMP= 0.0812); (C, right: di-18:1-BMP= 0.247; 2,2’ di- 18:1-BMP= 0.517; 2,3’ di-18:1-BMP= 0.098; 3,3’ di-18:1-BMP= 0.0005).

Nevertheless, SNX13 still bypassed NPC1 deficiency and this could have been due to the unexpected increased BMP that may be sufficient to facilitate cholesterol export from lysosomes.

To verify this SNX13 knockout dependent BMP increase, mass spectrometry was also carried out to quantify BMP isoforms for cells with and without SNX13, ±U18666A. di- 22:6 BMP was the predominant form in this cell type but all forms detected showed a striking increase in SNX13 siRNA treated cells. Moreover, similar (but slightly decreased) levels of total BMP were seen in samples treated with U18666A, suggesting that any changes detected by microscopy reflect an antibody-accessible pool that accumulates when NPC1 function is blocked during the 16h U18666A incubation time frame. Control experiments confirmed increased antibody-detected, endolysosomal BMP levels in U18666A-treated U2OS cells (Fig. S5 E, F). Altogether, these data demonstrate that SNX13 function is tightly connected to regulation of BMP levels in U2OS cells.

## DISCUSSION

We have presented here parallel screens to investigate the pathways that regulate cholesterol and BMP in cells. Our findings complement other previous screens related to cholesterol trafficking and highlight the strong, coordinate regulation of cholesterol and BMP. Despite the importance of BMP in cholesterol transport and other lipid degradative processes, little is known about how this lipid is synthesized or degraded. Strikingly, no single, specific metabolic enzyme revealed itself as a strong candidate for BMP biosynthesis. Rather, knockout of the mannose 6-phosphate receptor yielded the strongest phenotype. The simplest explanation is that BMP is upregulated in response to a severe defect in lysosomal enzyme delivery. Alternatively, BMP degradation is blocked.

Our complementary screens generated an enormous dataset that not only recapitulates many known trafficking, metabolic and transcriptional pathways and proteins that regulate intracellular cholesterol physiology, but importantly, it also revealed previously unrecognized roles for many genes and protein complexes that deserve further validation and characterization, opening new avenues for future investigation. One example is the identification of the PRC2 complex, whose subunits reduced cholesterol levels upon knockout only when NPC1 function was blocked (Fig. 5). Another example is the discovery of RP11-512M8.5, which encodes a protein of unknown function that shows high homology (>90%) to VPS33A, a HOPS/CORVET subunit. Consistently, RP11-512M8.5 behaved exactly the same as other uncovered HOPS/CORVET core subunits did: increasing or decreasing cholesterol in control or U18666A conditions, respectively (Fig. 3, Fig. S2).

In addition, our results also revealed unexpected consequences of deleting well-characterized players. For example, why does deletion of any of the ESCRT subunits consistently increase cholesterol under NPC1 inhibition conditions (Fig. 5, Fig. S2)? Depletion of ESCRT complexes has been shown to decrease, but not abolish, the number of intralumenal vesicles in multi-vesicular endosomes (Stuffers et al., 2009). In other work, depletion of the ESCRT-0 subunit, HRS but not other ESCRT subunits caused an NPC-like phenotype (Du et al., 2012). It is possible that this phenotype is only seen upon full knockout of ESCRTs carried out here. In the absence of ESCRT function, the alternative, ceramide-dependent intralumenal vesicle biogenesis pathway may be up-regulated (Stuffers et al., 2009; Trajkovic et al, 2008), perhaps generating intralumenal vesicles with higher cholesterol content. Further work is needed to resolve this question.

SNX13 was an especially interesting hit that when knocked out, could rescue lysosomal cholesterol accumulation in cells lacking NPC1 function, triggered either via NPC1 knockout or U18666A inhibition. Moreover, under these conditions, accessible cholesterol was detected at the cell surface compared with control cells, despite lack of NPC1 function. How might NPC1 function be bypassed? NPC1 bypass can be achieved by treatment of cells with cyclodextrin (Abi-Mosleh et al., 2009; Rosenbaum et al., 2010); in this case, it is likely that endocytosed cyclodextrin enables hydrophobic cholesterol to permeate the glycocalyx and gain access to the limiting lysosome membrane for eventual transport to the ER. Cyclodextrin may also trigger multivesicular endosome fusion with the cell surface, triggering cholesterol release.

Another example of NPC1 bypass was recently reported by Spiegel and colleagues (Newton et al., 2020). Remarkably, activation of sphingosine kinase was sufficient to drive cholesterol export from NPC1-deficient lysosomes. Sphingolipids are normally degraded by lysosomal sphingomyelinase, yielding sphingosine that is transported into the cytoplasm. There, phosphorylation by sphingosine kinase pulls overall sphingosine export by creating a substrate for a cytoplasmic degradative lyase; alternatively, sphingosine can be reutilized for ceramide synthesis in the ER. There are several possible explanations for this NPC1 bypass. First, increased sphingosine kinase could increase membrane contact sites (Höglinger et al., 2019; Meneses-Salas et al., 2020), thereby providing access of accumulated lysosomal cholesterol to an alternative cholesterol export route. It is also possible that NPC2 could deliver cholesterol directly to lumenal membranes that contain a less dense glycocalyx such as those of late endosomes (Cheruku et al., 2006; McCauliff et al., 2019) where LAMP proteins that contribute to the glycocalyx are less abundant (Li et al., 2016). In this case, enhanced sphingosine kinase could somehow enhance late endosome-lysosome fusion to permit such egress. Alternatively, sphingosine kinase activation could enhance multi-vesicular endosome exocytosis (Kajimoto et al., 2013), releasing accumulated cholesterol into the extracellular space.

In relation to our findings, SNX13 appears to be a negative regulator of cholesterol export from lysosomes such that export is enhanced in its absence. As an organelle tether (Henne et al., 2015), SNX13 may coordinate the association of membrane bound compartments to maintain appropriate and tightly regulated cholesterol levels in distinct cellular compartments, particularly in the ER. Further work will be needed to parse the differences in phenotypes observed for the related but distinct SNX13 and SNX14 proteins that both influence cholesterol and neutral lipid accumulation (Bryant et al., 2018; Datta et al., 2019, 2020).

Recently, Trinh et al. discovered a surprising role for PTDSS1, a phosphatidylserine (PS) synthase, in transport of cholesterol from the plasma membrane to the ER. PTDSS1 was also detected in our screens. In addition, the ER-localized scramblases, TMEM41B and VMP1, were reported to regulate normal distribution of cholesterol and phosphatidylserine (Li et al., 2021). These two genes were also identified as hits in our cholesterol screens, both increasing cholesterol only under NPC1-inhibition conditions, similar to PTDSS1 (Fig. S2, S3). Knockout of either scramblase resulted in cholesterol plasma membrane re-distribution, a phenotype reminiscent of that observed upon PTDSS1 knockout (Trinh et al., 2020). In our screens, SNX13 was uncovered as a cholesterol-decreasing hit (Fig. 2, Fig. 3); this is probably because SNX13 plays a broader role in the homeostatic regulation of lipids, controlling for example the levels of BMP and fatty acids in the cell (Figs. 8, 9). Note that SNX13 depletion enables cholesterol to reach the plasma membrane despite loss of NPC1 function; this likely differs from plasma membrane accumulation seen upon knockout of PTDSS1 and the scramblases, as levels did not appear to represent accumulation.

Importantly, PS is likely important for the formation of ER-plasma membrane junctions by ER-associated GRAMD1/Aster proteins that bind anionic lipids and transfer cholesterol via membrane proximal START or START like domains (Naito et al., 2019; Sandhu et al., 2018). Like GRAMD/Aster proteins, SNX13 is an ER-anchored membrane protein with lipid binding motifs and organelle tethering potential; its Drosophila Snz homolog binds phosphoinositides as well as phosphatidylserine (Ugrankar et al., 2019). However, unlike GRAMD/Aster proteins, SNX13 does not contain an obvious sterol binding domain. It is possible that SNX13 links the ER to both late endosomes and the plasma membrane, and future experiments will explore the possibility that PS binding is important for its cholesterol regulation activities. Finally, paradoxically, NPC loss has been reported to increase membrane contact sites (Lim et al., 2019), yet certain of these can rescue the cholesterol accumulation phenotype (Höglinger et al., 2019; see also Meneses-Salas et al., 2020). Thus, not all membrane contact sites are created equal and their regulation and specificity will be important to address in future work.

## Materials and Methods

### Cell culture, antibodies and other reagents

U2OS, HeLa and RPE cells were obtained from ATCC and grown in Dulbecco’s modified Eagle’s media (DMEM) containing 10% fetal bovine serum, 2 mM L-glutamine, and penicillin (100 U/ml)/ streptomycin (100 μg/ml). Cas9-expressing K562 cells (Lu et al., 2018) were grown in RPMI 1640 Medium supplemented with 10% fetal bovine serum, 2 mM L-glutamine, 1 mM sodium pyruvate and penicillin (100 U/ml)/streptomycin (100 μg/ml). All cell lines were cultured at 37°C with 5% CO_2_. The NPC1 CRISPR knockout HeLa cell line was previously generated (Saha et al., 2020). Primary antibodies diluted in PBS- 1%BSA (for immunofluorescence) or 5% skim milk (for immunoblotting) were monoclonal Anti-LBPA (BMP) clone 6C4 1:1000 (Millipore, Cat# MABT837), rabbit polyclonal anti-LAMP2 1:500 (Invitrogen, Cat# PA1-655), mouse monoclonal anti-GST 1:1000 (Cell Signaling, Cat# 2624S), rabbit polyclonal anti-SNX13 1:1000 (Abcepta, Cat# AP12244b), mouse monoclonal anti-Tubulin 1:10000 (Sigma-Aldrich, Cat# T5168), rabbit monoclonal anti-HA Tag (Cell Signaling, Cat# 3724S). U18666A (Sigma-aldrich; Cat# U3633) was used at 1µM for 16h. Oleic acid (Sigma-Aldrich; Cat# O1008,) was conjugated with fatty-acid free BSA (Sigma-Aldrich; A8806) at a 6:1 molar ratio and used at 0.5 mM for 16h. The ACAT pharmacological inhibitor Sandoz 58-035 (Sigma-Aldrich; S9318) was used at 20 µg/mL for 16h.

### Plasmids and transfections

Plasmids encoding SNX13-GFP and SNX14-GFP were a gift from Mike Henne (UT Sothwestern). The CFP-VAPA construct was a gift from Clare Futter (University College London). The WT-SNX13-2XHA plasmid was prepared as follows: First, the BioID2 insert from a target MCS-BioID2-HA plasmid (Addgene, Cat# 74224) was removed by restriction digestion at BspEI and HindIII sites and replaced with a 2XHA oligo sequence flanked by the same restriction sites. Next, the WT-SNX13 ORF sequence from the SNX13-GFP plasmid was shuttled into the NheI and BspEI restriction sites of the previously modified MCS- BioID2-HA. The SNX13 truncated constructs were cloned by Gibson assembly using WT- SNX13-2XHA as template (see primers in Supplementary Table 1). Transfections of U2OS cells plated on coverslips were carried out using GenJet Plus Reagent (SigmaGen Laboratories; Cat# SL10050) according to the manufacturer. For siRNA-mediated knockdown experiments, cells were transfected with siRNAs targeting human SNX13 (5’- CAGAAAGGCUCAACAGAAAUU-3’) or SNX14 (5’-GGAUGAAAGUAUUGACAAAUU-3’) using Lipofectamine RNAiMax (Invitrogen, Cat#13778-075) according to the manufacturer. Studies were conducted 48-72 h after siRNA transfection. A Scrambled siRNA was used as negative control (Ambion, Cat# AM4635).

### Quantitative real-time PCR

Total RNA from SNX14 siRNA- or control siRNA-treated cells was extracted using QIAzol (Qiagen, Cat# 79306) as per the manufacturer. 1 μg isolated RNA was reverse-transcribed using High Capacity cDNA Reverse Transcription Kit (Applied Bioscience, Cat# 4368814). qRT-PCR was performed using SYBRGreen-based technology GoTaq® qPCR master-mix (Promega, Cat# A600A). Specific SNX14 primers, and HPRT1 primers (reference gene) were used (see Table 1). PCRs were run in a LightCycler 96® Real-time PCR system (Roche). Transcript levels relative to HPRT1 were calculated using the ΔCt method.

### Flow cytometry analysis of Cholesterol and BMP

K562 cells were treated ± 1µM U18666A. After 16h cells were harvested and washed before fixation in 2% paraformaldehyde, then permeabilized in 0.1% Saponin and stained with PFO*-Alexa647 or anti-BMP and Alexa Fluor 647 goat anti–mouse antibodies (Invitrogen). Quantification of cholesterol and BMP total fluorescence was performed using a BD Accuri C6 flow cytometer and data were analyzed using Flowjo software (Tree Star).

### PFO* purification

Recombinant PFO* was purified as previously described (Li, Lee and Pfeffer, 2017). Briefly, expression was induced in Rosetta 2 cells with 1 mM isopropyl β-p-thio- galactopyranside for 4 h at 37 °C. Cells were resuspended in buffer A [PBS, 10% (vol/vol) glycerol, protease inhibitors] and lysed using an Emulsiflex C-5 homogenizer (Avestin). Clarified lysate was incubated with Ni-NTA agarose (Qiagen, Cat# 30210) for 1 h at 4 °C and after washing with buffer A + 50 mM imidazole, bound protein was eluted with buffer A + 300 mM imidazole. The eluate was concentrated with an Amicon Ultra 10-kDa cutoff centrifugal filter (Millipore, Cat# UFC901024) and then exchanged into buffer A + 1 mm EDTA. PFO* was directly conjugated to NHS ester Alexa Fluor 647 dye as per the manufacturer (Life Technologies). For immunofluorescence analysis of cholesterol, GST-PFO* was used (Meneses-Salas et al. 2020).

### CRISPR/Cas9-bEXOmiR screens in K562 cells

The Genome-wide K562 CRISPR knockout library line was generated as previously described (Morgens et al., 2017). Briefly, a whole-genome library of exon-targeting sgRNAs (10 sgRNA per gene; Morgens et al., 2017) was synthesized and cloned into a lentivirus vector (Addgene, Cat# 89359) which together with third-generation lentiviral packaging plasmids (pVSVG, pRSV and pMDL) were transfected into HEK293T cells to generate lentiviral particles. Then ∼300 million Cas9-expressing K562 cells were infected at low MOI (<1). Transduced cells were selected and expanded in puromycin-supplemented media over 5–7 days before conducting experiments 10 days post-infection. All screens were performed as independent replicates. Two independent screens were performed for each lipid (cholesterol and BMP) and condition (±U18666A). For each screen, 600 million cells were stained. 16h before staining, cells were treated with 1µM U18666A or vehicle (control). Next day, cells were first pelleted and washed twice in cold PBS followed by fixation in 2% paraformaldehyde-PBS for 30 min at 4°C. Cells were then washed twice in PBS and permeabilized/blocked in 0.1%Saponin-1%BSA-PBS for 10 min. Cells were then incubated with 10 µg/ml Alexa-647 labeled Perfringolysin O* (Li et al., 2017) for 45 min in the cold. Alternatively, for BMP screens, staining was performed with mouse anti-BMP antibody for 1 h at 4°C, followed by 1 h incubation with Alexa 647-conjugated secondary antibody (Life Technologies) used at 1:2,000. Finally, cells were washed once in cold PBS and then kept in 15 mL PBS-0.5%BSA at 4°C for 16h before sorting. Next day, cells were separated into 10% high or 10% low PFO/BMP fluorescence populations by sorting on a BD FACSAria^TM^II. Around 20 million cells were recovered from each gated population. Sorted cells were then sedimented by centrifugation, and the cell pellet was frozen at -80°C before genomic DNA isolation. Approximately 200 million unsorted cells (1000x coverage per library element) were saved for screen data normalization. Genomic DNA was extracted using Qiagen Blood Midi or Maxi kits (Qiagen, Cat# 51183; 51194) for sorted or unsorted cells respectively, as per the manufacturer. To prepare sequencing libraries, the sgRNAs sequences were PCR-amplified from genomic DNA and the number of PCR reactions was scaled to use 40-60 µg isolated genomic DNA. Each of these 1st PCR reactions contained: 10 µg of genomic DNA, 2 µL Herculase II Fusion DNA polymerase (Agilent Technologies, Cat# 600677), 20 µL 5X Herculase buffer, 1 µL 100 mM dNTPs, 1 µL 100 µM oMCB_1562 (forward primer), 1 µL 100 µM oMCB 1563 (reverse primer) and water to adjust the volume to 100 µL. PCR amplification was conducted as follows: 1x 98°C/2 min, 18x 98°C/30 s, 59.1°C/30 s, 72°C/45 s, 1x 72°C/3 min. 1st PCR reactions from each sample were pooled and then 2nd PCR reactions were set up for each sample as follows: 5 µL from 1st PCR pooled amplicons, 2 µL Herculase II Fusion DNA polymerase, 20 µL 5X Herculase buffer, 2 µL of 100 mM dNTPs, 0.8 µL 100 µM oMCB_1439 (forward primer), 0.8 µL 100 µM of barcoded CRISPR KO reverse primer, and 69.4 µL H_2_O. PCR protocol for this 2nd PCR reaction was: 1x 98°C/2 min, 20x 98°C/30 s, 59.1°C/30 s, 72°C/45 s, 1x 72°C/3 min. Finally, 50 µL of 2nd PCR reaction was separated by running on a 2% TBE-agarose gel. The PCR products were excised and purified using a QIAquick Gel Extraction Kit (Qiagen, Cat# 28704) according to the manufacturer’s instructions. The sgRNA libraries were analyzed by deep sequencing on an Illumina NextSeq 500 using a custom sequencing primer (oMCB_1672), with ∼40 million reads per condition (∼200x coverage per library element). Computational analysis and comparison of sgRNA composition of sorted versus unsorted cells were performed using casTLE v.1.0 (https://bitbucket.org/dmorgens/castle) as previously described (Lu et al., 2018; Morgens et al., 2016). Briefly, sgRNA distribution was compared between the sorted and unsorted cell samples and sgRNA enrichments were calculated as log ratios between sorted and unsorted cells. A maximum likelihood estimator was used to estimate the phenotypic effect size for each gene and the log-likelihood ratio (confidence score) by comparing the distribution of the 10 different sgRNAs targeting each gene to the distribution of negative control sgRNAs. P values were determined by permuting the gene-targeting sgRNAs in the screen and comparing to the distribution of negative controls using casTLE. For genome-wide cholesterol screens, we used a threshold of 5% FDR (calculated using the Benjamini-Hochberg procedure) to define hits. For the BMP screens, the top 100 ranked genes in the analysis were considered as hits. Because cholesterol is functionally linked to BMP (Chevallier et al 2008), despite lower statistical significance for some of the BMP screen hits, we found that a significant number of genes that passed this cutoff overlapped with those identified as hits in the cholesterol screens. See Supplementary Table 1 for complete genome-wide screen datasets.

### Immunofluorescence

To visualize endolysosomes in suspension K562 cells (Figure 1B-D), cells were attached to glass coverslips using a cytospin (Shandon) at 800 rpm for 5 min. Cells were fixed with 3.7% (v/v) paraformaldehyde for 15 min, washed and permeabilized/blocked with 0.1% saponin/1% BSA-PBS except for cells labeled with GST-PFO*, which were permeabilized for 3 min in 0.1% Triton X-100 and blocked with 1% BSA in PBS. Primary antibodies were diluted in PBS-1%BSA and incubated for 1h at RT. Highly cross-adsorbed H+L secondary antibodies (Life Technologies) conjugated to Alexa Fluor 488, 568, or 647 were used at 1:2,000 in PBS-1%BSA and incubated at RT for 45 min. Nuclei were stained using 0.1μg/ml DAPI (Sigma-Aldrich,Cat# D9542) and coverslips were mounted on glass slides with Mowiol. Microscopy images were acquired using a Zeiss LSM880 laser scanning spectral confocal microscope (Carl Zeiss, Germany) equipped with an Axio Observer 7 inverted microscope, blue diode (405nm), Argon (488nm), diode pumped solid state (561nm) and HeNe (633nm) lasers and a Plan Apochromat 63x numerical aperture (NA) 1.4 oil-immersion objective lens was used. DAPI, Alexa 488, Alexa Fluor 555 and Alexa Fluor 647 images were acquired sequentially using 405, 488, 561and 633 laser lines, AOBS (Acoustic Optical Beam Splitter) as beam splitter and emission detection ranges 415- 480, 500-550 nm, 571-625 nm and 643-680nm respectively. Confocal pinhole was set at 1 Airy units. All images were acquired in a 1024 x 1024 pixel format. In some experiments (Figure 1B-D, Supp. Figure 4B), images were obtained using Metamorph software with a spinning disk confocal microscope (Yokogawa) with an electron multiplying charge-coupled device (EMCCD) camera (Andor) and a 100x 1.4 NA oil-immersion objective. Typical exposure times of 100–300 ms were used. 3D-rendered images (Supplementary Figure 4E) were generated using IMARIS software (Bitplane AG). All image quantifications were performed using CellProfiler (Carpenter et al., 2006).

### Immunoblotting

Cells were lysed in lysis buffer (50 mM HEPES, 150 mM KCl, 1% Triton X- 100, 5 mM MgCl_2_, pH 7.4) supplemented with a protease/phosphatase inhibitor cocktail (1mM Na_3_VO_4_, 10 mM NaF, 1 mM PMSF, 10 μg/ml leupeptin and 10 μg/ml aprotinin). Lysates were boiled in 1x sample buffer, resolved on SDS-PAGE and transferred onto nitrocellulose membranes (Bio-Rad, Cat# 1620115) using a Bio-Rad Trans-blot system. Membranes were blocked with 5% skim milk in Tris-buffered saline with Tween-20 for 60 min at RT. Primary antibodies were diluted in blocking buffer and incubated either 1 h at RT or overnight at 4°C. HRP-conjugated secondary antibodies (Bio-Rad or Abcam) diluted in blocking buffer at 1:5,000 were incubated for 60 min at RT and developed using enhanced chemiluminescence EZ-ECL (Biological Industries). Blots were imaged using an ImageQuant LAS 4000 system (GE Healthcare) and quantified using ImageJ software.

### Thin Layer chromatography

Total Lipids were extracted from control or SNX13-depleted cells as follows. Briefly, cells were washed with PBS and resuspended in 1 volume of Methanol:Chloroform (1:2). Next 1/2 volume chloroform and 1/2 volume of H_2_O were added and tubes were centrifuged at 5000 RPM for 5 min. The organic phase was collected, dried, resuspended with chloroform and spotted on on TLC silica gel 60 plates (Millipore, Cat# 1055530001), and dried for several minutes. Plates were run in a solvent system of hexane/diethyl ether/acetic acid (70:30:1) until 3/4 of the total length of the plate was reached. Lipids were stained using a phosphomolybdic acid (ACROS organics, Cat# 206385000)-ethanol solution.

### Bis(monoacylglycero)phosphate (BMP) lipidomics

Targeted high resolution UPLC-MS/MS was used to accurately quantitate the three geometrical isoforms (2,2’-, 2,3’-, and 3,3’-) of di-22:6- BMP and di-18:1-BMP in control or SNX13 siRNA-treated cells ±U18666A. Lipidomics analyses were conducted by Nextcea, Inc. (Woburn, MA) as previously described (Liu et al., 2014) using a SCIEX TripleTOF 6600 mass spectrometer equipped with an IonDrive Turbo V source (SCIEXm Framingham, MA). Standard curves were prepared using authentic BMP reference standards.

### Functional Network analysis

Top hits from both cholesterol and BMP screens were collectively queried in STRING (http://string-db.org/) to search for experimentally-confirmed interactions with a high confidence score (0.7). The resulting STRING graphics file reporting interactions was then manually curated using Adobe Illustrator software (Adobe) to create a visually comprehensive functional interactome. Interactions were represented as edges whose thickness was proportional to a calculated score derived from STRING analysis. Additional relevant interactions not reported by STRING but confirmed by the BioGRID database (http://thebiogrid.org/) were also displayed as green edges. The final interactome map was further manually curated to include additional screen hits (nodes) that despite not being reported to interact, could still be clustered in a common functional category revealed by the screens performed.

### Statistics

Results are expressed as mean ± SEM unless otherwise specified. Means were compared using Student’s *t* test where two experimental conditions were compared. When three or more experimental conditions were compared statistical significance was assessed via multiple *t* test by Holm-Sidak method with α = 0.05 or two-way ANOVA with Tukey’s post hoc test using Graph Pad Prism 9. Two-tailed P values < 0.05 were considered statistically significant (Lord et al., 2020).

## Supplemental material

The supplemental material includes 5 figures and one Excel file providing all of the data derived from the screens described herein.

Supplemental Figure 1. Bivariate analyses comparing hits in relation to changes in cholesterol and BMP.

Supplemental Figure 2. Subcellular localization of screen hits and their phenotypes.

Supplemental Figure 3. Lipid metabolic pathways revealed in these screens.

Supplemental Figure 4. SNX13 is an ER resident protein that associates with lipid droplets via its C-terminus.

Supplemental Figure 5. SNX13 depletion redistributes cholesterol to the cell surface of HeLa cells in the absence of NPC1 function.

Supplemental Table 1. Oligonucleotide sequences used in this study and all screen results as an Excel file.

## Supporting information

Supplemental Table 1 Screen Data

## Acknowledgments

This research was funded by grants to S.R.P from the NHLBI (5R01HL134991-04) and the Ara Parseghian Medical Research Foundation/Notre Dame University. C.E. was funded by grants BFU2015-66785-P from the Spanish Ministerio de Economía y Competitividad and AR0RM005 from the University of Barcelona (Spain).The authors declare no competing financial interests.

## Author contributions

All experiments were carried out by A.L. F.H. carried out the mass spectrometric analysis of BMP. C.E. provided materials and advice and S.R.P oversaw the project, obtained research funding and A.L. and S.R.P. wrote and edited the manuscript.

---

**Supplemental Figure 1.**
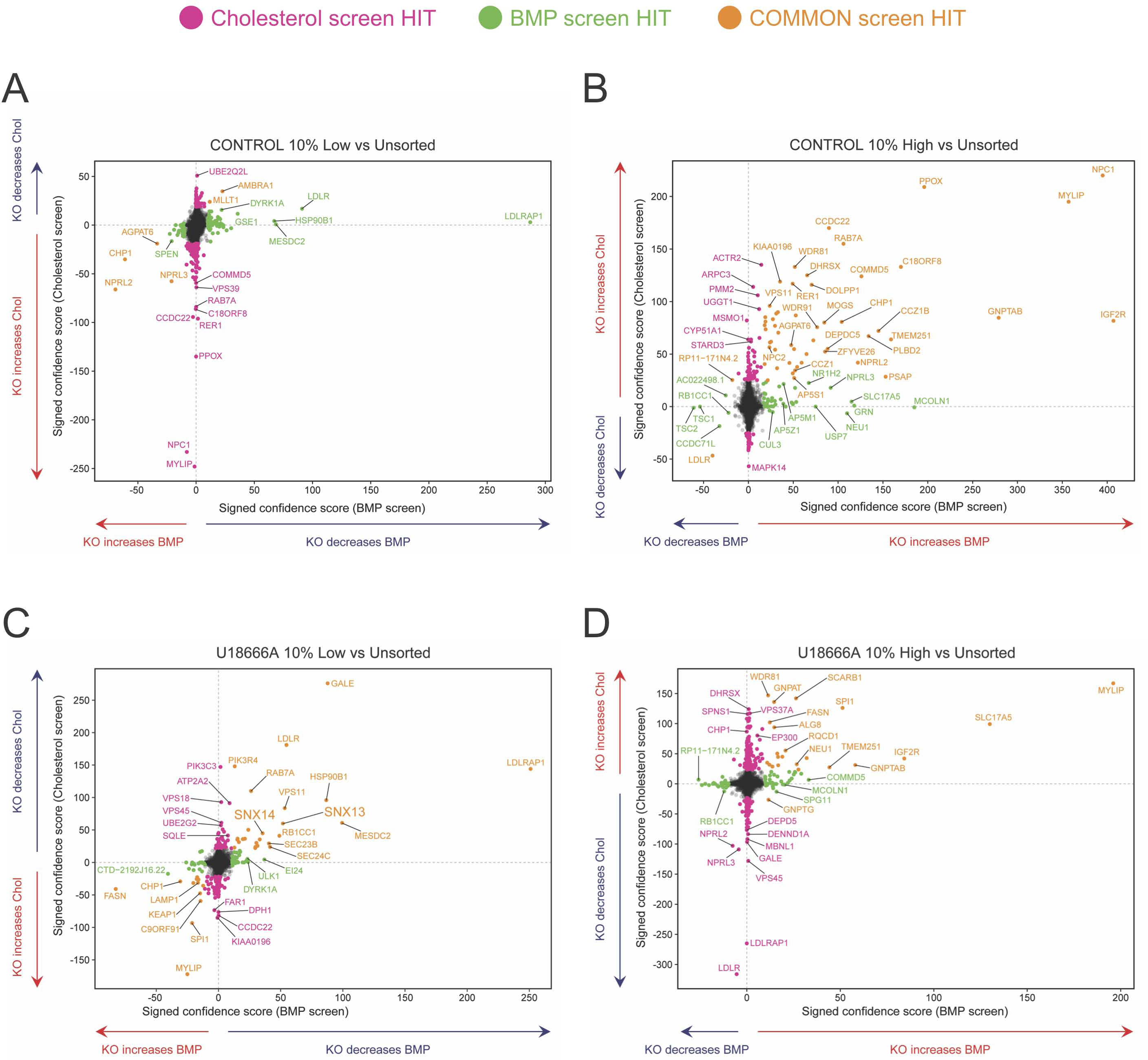
Bivariate analyses comparing hits in relation to changes in cholesterol and BMP. Indicated genes are presented in relation to their phenotypes determined by PFO* and BMP detection and flow cytometry, presented as signed confidence scores. Genes that appeared in both analyses are shown in goldenrod; those seen only in the cholesterol or BMP screens are shown in pink or green, respectively. The panels represent hits discovered in (A), 10% low versus unsorted; (B), 10% high versus unsorted; (C and D) same as in (A and B) but in the presence of U18666A.

**Supplemental Figure 2.**
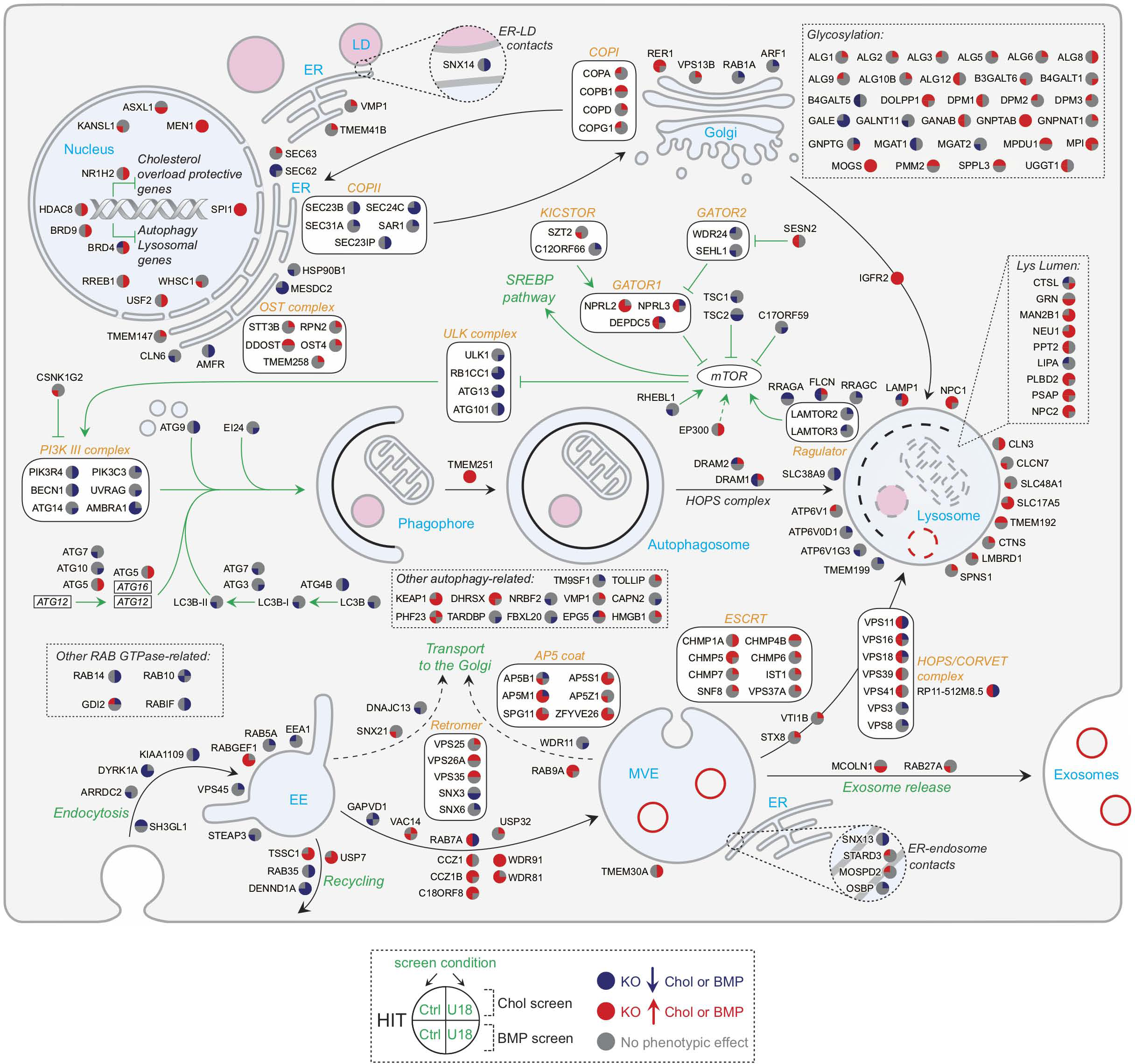
Subcellular localization of screen hits and their phenotypes. Selected top hits are annotated in colored circles that show increases (red) or decreases (blue) in cholesterol upper left, BMP, lower left, cholesterol +U18666A (upper right) or BMP + U18666A (lower right) as indicated. Black arrows indicate trafficking routes. Green arrows indicate activation; inhibition signs are also displayed in green. ER= Endoplasmic reticulum, LD= Lipid droplet, EE= Early endosome, MVE= Multivesicular endosome.

**Supplemental Figure 3.**
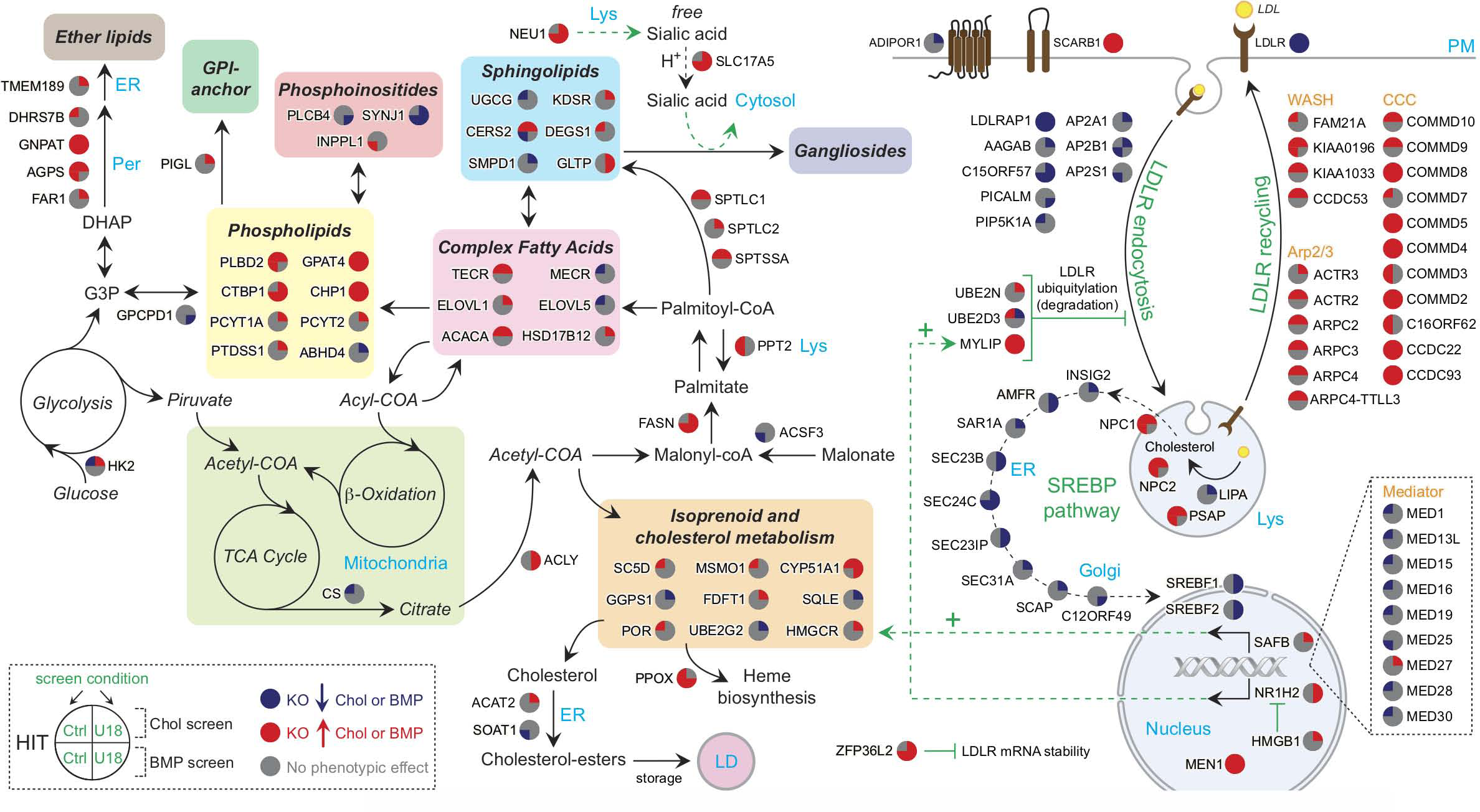
Lipid metabolic pathways revealed in these screens. Genes are annotated in colored circles that show increases (red) or decreases (blue) in cholesterol upper left, BMP, lower left, cholesterol +U18666A (upper right) or BMP + U18666A (lower right) as indicated. Black arrows indicate trafficking, metabolic routes and enzymatic reactions. Green dotted arrows indicate activation of gene expression of the indicated gene/s. Inhibition signs are displayed in green. ER= Endoplasmic reticulum, Per= Peroxisome, PM= Plasma membrane, LD= Lipid droplet, Lys= Lysosome.

**Supplemental Figure 4.**
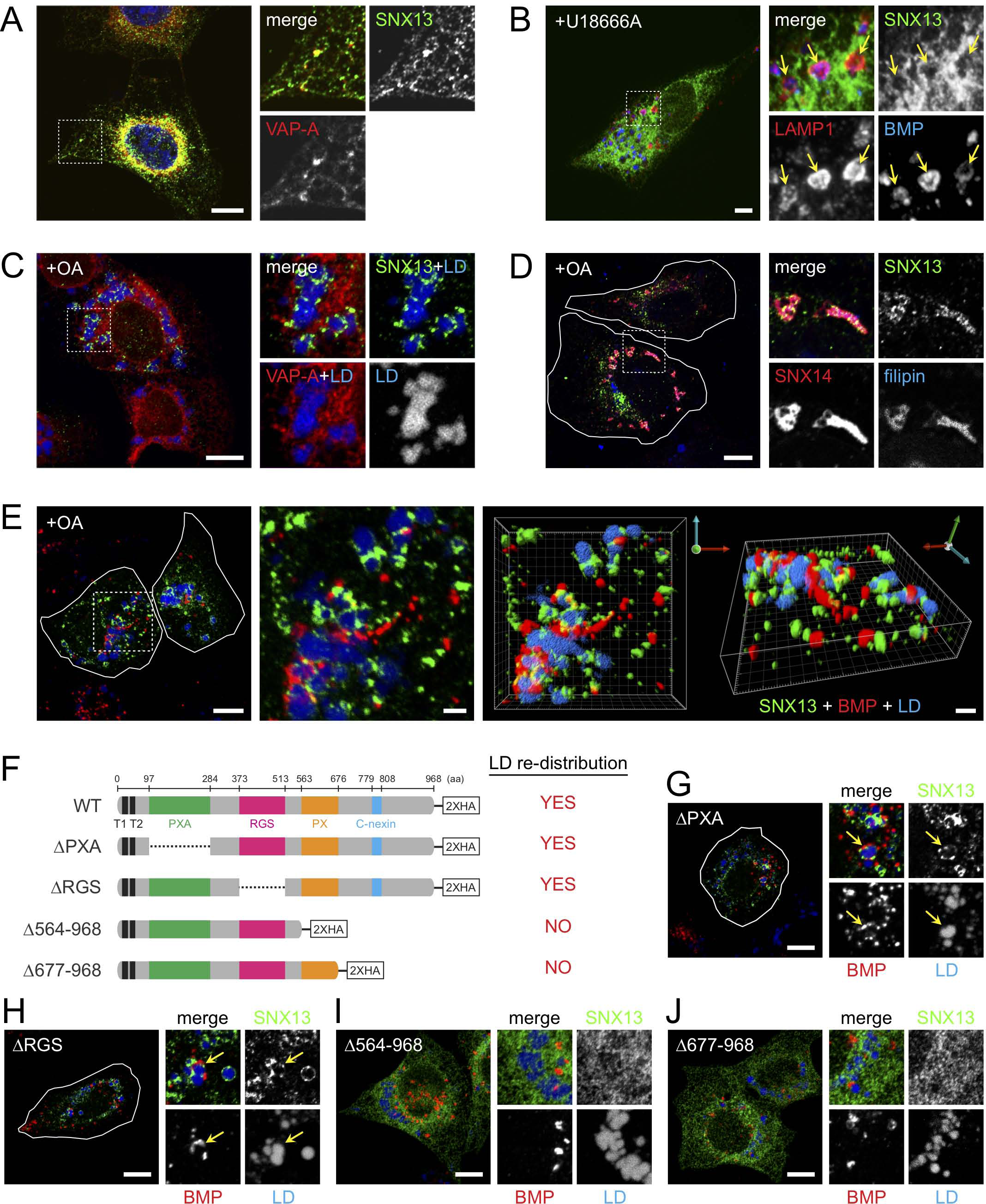
SNX13 is an ER resident protein that associates with lipid droplets via its C-terminus. (A) Immunofluorescence microscopy of U2OS cells expressing SNX13-HA and VAP-A-CFP. Proteins were detected with anti-HA antibodies or CFP fluorescence as indicated; scale bar, 10µm. (B) RPE cell expressing SNX13-GFP, treated with U18666A for 16 hours. LAMP1 and BMP were detected using specific antibodies. Scale bar, 10µm. (C, D) U2OS cells as in A, treated with oleic acid overnight. Lipid droplets (LDs) were detected using LipidTOX; cholesterol was detected using filipin. VAP-A-CFP and SNX14-GFP were detected using their intrinsic fluorescence. Shown in small boxes are enlargements of the boxed areas shown at left; scale bars, 10µm. (E) Cells as in (A) were visualized by confocal microscopy; green, SNX13; red, BMP; blue, LDs. Scale bar, 10µm except for enlarged inset and 3D rendering, 2µm. (F) Schematic analysis of SNX13 constructs. Colored regions indicate domain organization as indicated. LD localization is summarized. (G-J), localizations of constructs indicated in (F) Cells were labeled as in (E) with the indicated markers. Scale bars, 10µm.

**Supplemental Figure 5.**
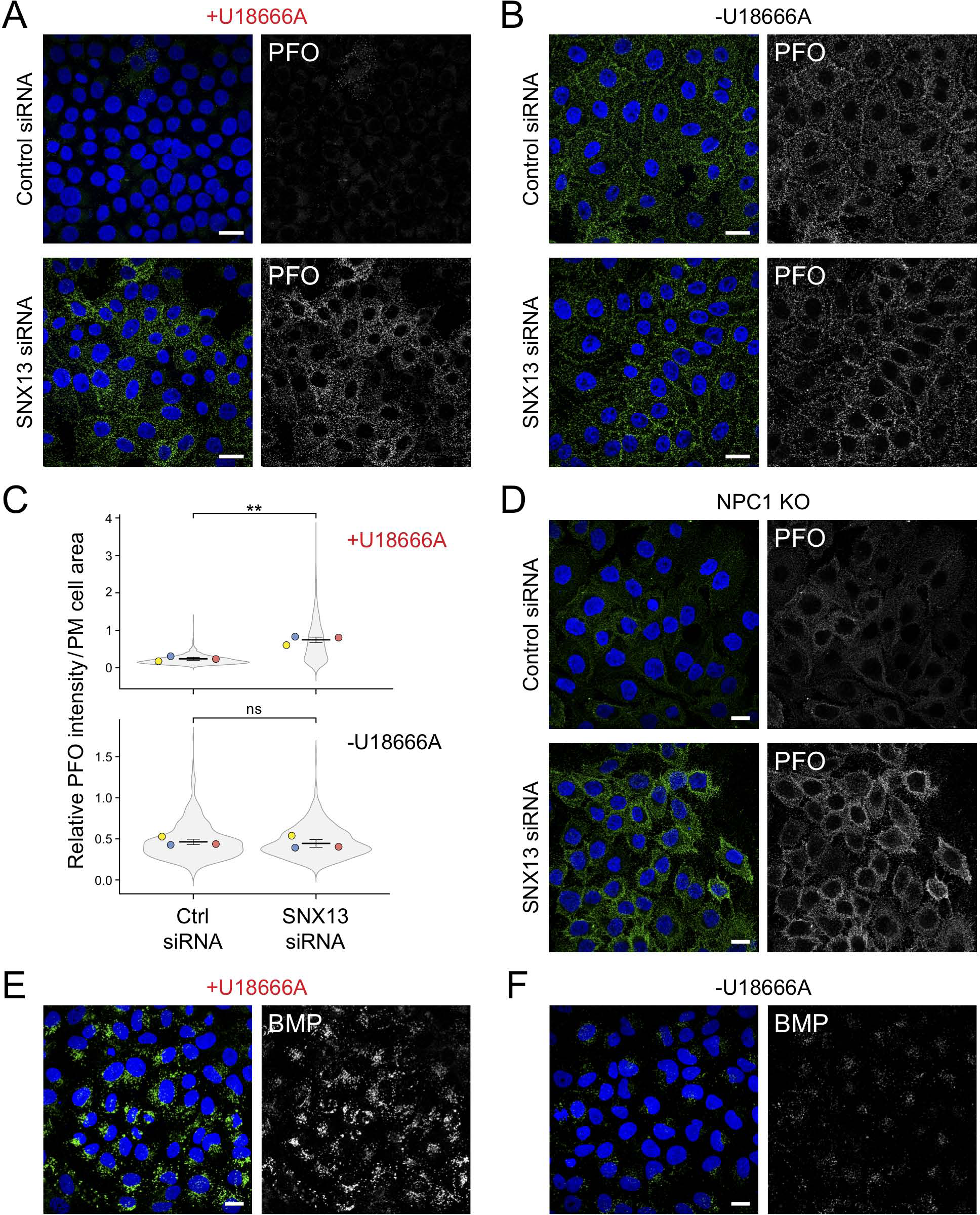
SNX13 depletion redistributes cholesterol to the cell surface of HeLa cells in the absence of NPC1 function. (A and B) Immunofluorescence microscopy of HeLa cells treated with U18666A for 16 hours (A) or left untreated (B), and labeled in the absence of detergent permeabilization with with GST-PFO* detected using anti-GST primary antibodies, 72 hours after transfection with the indicated siRNAs. (C and D) Quantitation of PFO* staining as a function of plasma membrane cell area in U18666A-treated (C) and untreated (D) cells, determined using CellProfiler. Colored dots reflect means from independent experiments; >830 cells analyzed in each condition, significance was determined by unpaired *t* test; **, P < 0.01. (D) Immunofluorescence microscopy of NPC1-knockout HeLa cells stained in the absence of detergent permeabilization with GST-PFO detected by anti-GST primary antibodies. (E and F) U2OS cells treated with U18666A for 16 hours (E) or left untreated (F), labeled with anti-BMP antibodies and imaged by confocal microscopy. Scale bars, 20µm. Actual P values: (C, top: Ctrl siRNA+U18 vs SNX13 siRNA+U18= 0.0017); (C, bottom: Ctrl siRNA vs SNX13 siRNA= 0.377).

## Notes

### Competing Interest Statement

The authors have declared no competing interest.

## References

Abi-Mosleh L, Infante RE, Radhakrishnan A, Goldstein JL, and Brown MS. 2009. Cyclodextrin overcomes deficient lysosome-to-endoplasmic reticulum transport of cholesterol in Niemann-Pick type C cells. Proc. Natl. Acad. Sci. U S A 106:19316–21.

Adachi, S., Homoto, M., Tanaka, R., et al. 2014. ZFP36L1 and ZFP36L2 control LDLR mRNA stability via the ERK-RSK pathway. Nucleic Acids Res. 42:10037–10049.

Aregger, M., Lawson, K. A., Billmann, M., Costanzo, M., Tong, A., Chan, K., Rahman, M., Brown, K. R., Ross, C., Usaj, M., Nedyalkova, L., Sizova, O., Habsid, A., Pawling, J., Lin, Z. Y., Abdouni, H., Wong, C. J., Weiss, A., Mero, P., Dennis, J. W., et al. 2020. Systematic mapping of genetic interactions for de novo fatty acid synthesis identifies C12orf49 as a regulator of lipid metabolism. Nat Metab. 2:499–513.

Ballabio A, and Bonifacino JS. 2020. Lysosomes as dynamic regulators of cell and organismal homeostasis. Nat. Rev. Mol. Cell Biol. 21:101–118.

Bayraktar, E. C., La, K., Karpman, K., Unlu, G., Ozerdem, C., Ritter, D. J., Alwaseem, H., Molina, H., Hoffmann, H. H., Millner, A., Atilla-Gokcumen, G. E., Gamazon, E. R., Rushing, A. R., Knapik, E. W., Basu, S., and Birsoy, K. 2020. Metabolic coessentiality mapping identifies C12orf49 as a regulator of SREBP processing and cholesterol metabolism. Nat Metab. 2:487– 498.

Bryant D, Liu Y, Datta S, et al. 2018. SNX14 mutations affect endoplasmic reticulum-associated neutral lipid metabolism in autosomal recessive spinocerebellar ataxia 20. Hum Mol Genet. 27:1927–1940.

Carpenter AE, Jones TR, Lamprecht MR, Clarke C, Kang IH, Friman O, Guertin DA, Chang JH, Lindquist RA, Moffat J, Golland P, Sabatini DM. 2006. CellProfiler: image analysis software for identifying and quantifying cell phenotypes. Genome Biology 7:R100.

Casanova, J. E., and Winckler, B. 2017. A new Rab7 effector controls phosphoinositide conversion in endosome maturation. J Cell Biol. 216:2995–2997.

Cerikan, B., Shaheen, R., Colo, G. P. et al. 2016. Cell-Intrinsic Adaptation Arising from Chronic Ablation of a Key Rho GTPase Regulator. Dev Cell 39: 28–43.

Cheruku SR, Xu Z, Dutia R, Lobel P, Storch J. 2006. Mechanism of cholesterol transfer from the Niemann-Pick type C2 protein to model membranes supports a role in lysosomal cholesterol transport. J Biol Chem. 281(42):31594–604.

Chevallier J, Chamoun Z, Jiang G, Prestwich G, Sakai N, Matile S, Parton RG, and Gruenberg J. 2008. Lysobisphosphatidic acid controls endosomal cholesterol levels. J. Biol. Chem. 283:27871–27880.

Chu BB, Liao YC, Qi W, Xie C, Du X, Wang J, Yang H, Miao HH, Li BL, and Song BL. 2015. Cholesterol transport through lysosome-peroxisome membrane contacts. Cell 161:291–306,

Das A, Goldstein JL, Anderson DD, Brown MS, Radhakrishnan A. 2013. Use of mutant 125I- perfringolysin O to probe transport and organization of cholesterol in membranes of animal cells. Proc Natl Acad Sci U S A. 110(26):10580–5.

Das A, Brown MS, Anderson DD, Goldstein JL, and Radhakrishnan A. 2014. Three pools of plasma membrane cholesterol and their relation to cholesterol homeostasis. Elife. 2014;3:e02882.

Datta S, Bowerman J, Hariri H, Ugrankar R, Eckert KM, Corley C, Vale G, McDonald JG, Henne WM. 2020. Snx14 proximity labeling reveals a role in saturated fatty acid metabolism and ER homeostasis defective in SCAR20 disease. Proc Natl Acad Sci U S A. 117(52):33282–94.

Datta S, Liu Y, Hariri H, Bowerman J, Henne WM. 2019. Cerebellar ataxia disease-associated Snx14 promotes lipid droplet growth at ER-droplet contacts. J Cell Biol. 218(4):1335–1351.

Davis OB, Shin HR, Lim CY, Wu EY, Kukurugya M, Maher CF, Perera RM, Ordonez MP, and Zoncu R. 2021. NPC1-mTORC1 Signaling Couples Cholesterol Sensing to Organelle Homeostasis and Is a Targetable Pathway in Niemann-Pick Type C. Dev. Cell 56:260–276.e7.

Diofano, F., Weinmann, K., Schneider, I. et al. 2020. Genetic compensation prevents myopathy and heart failure in an in vivo model of Bag3 deficiency. PLoS Genet. 16: e1009088.

Du, X., Kazim, A. S., Brown, A. J., et al. 2012. An essential role of Hrs/Vps27 in endosomal cholesterol trafficking. Cell Rep. 1:29–35.

Gruenberg J. 2020. Life in the lumen: The multivesicular endosome. Traffic 21:76–93.

Henne WM, Zhu L, Balogi Z, Stefan C, Pleiss JA, and Emr SD. 2015. Mdm1/Snx13 is a novel ER-endolysosomal interorganelle tethering protein. J. Cell Biol. 210:541–551.

Höglinger D, Burgoyne T, Sanchez-Heras E, et al. 2019. NPC1 regulates ER contacts with endocytic organelles to mediate cholesterol egress. Nat. Commun. 10:4276.

Infante RE, and Radhakrishnan A. 2017. Continuous transport of a small fraction of plasma membrane cholesterol to endoplasmic reticulum regulates total cellular cholesterol. Elife. 2017;6:e25466.

Kajimoto T, Okada T, Miya S, Zhang L, and Nakamura S. 2013. Ongoing activation of sphingosine 1-phosphate receptors mediates maturation of exosomal multivesicular endosomes. Nat. Commun. 4:2712.

Kolter, T., and Sandhoff, K. 2005. Principles of lysosomal membrane digestion: stimulation of sphingolipid degradation by sphingolipid activator proteins and anionic lysosomal lipids. Annu Rev Cell Dev Biol 21:81–103.

Li, J., Deffieu, M. S., Lee, P. L., Saha, P., and Pfeffer, S. R. 2015. Glycosylation inhibition reduces cholesterol accumulation in NPC1 protein-deficient cells. Proc Natl Acad Sci U S A. 112(48):14876–81.

Li, J., Lee, P. L., and Pfeffer, S. R. 2017. Quantitative measurement of cholesterol in cell populations using flow cytometry and fluorescent perfringolysin o. Methods Mol Biol. 1583:85– 95.

Li, Y. E., Wang, Y., Du, X., Zhang, T., Mak, H. Y., Hancock, S. E., McEwen, H., Pandzic, E., Whan, R. M., Aw, Y. C., Lukmantara, I. E., Yuan, Y., Dong, X., Don, A., Turner, N., Qi, S., and Yang 2021. TMEM41B and VMP1 are scramblases and regulate the distribution of cholesterol and phosphatidylserine. J Cell Biol. 220:e202103105.

Lim, C. Y., Davis, O. B., Shin, H. R., Zhang, J., Berdan, C. A., Jiang, X., Counihan, J. L., Ory, D. S., Nomura, D. K., and Zoncu, R. 2019. ER-lysosome contacts enable cholesterol sensing by mTORC1 and drive aberrant growth signalling in Niemann-Pick type C. Nat. Cell Biol. 21:1206–1218.

Liu, N., Tengstrand, E. A., Chourb, L., and Hsieh, F. Y. 2014. Di-22:6-bis(monoacyl- glycerol)phosphate: A clinical biomarker of drug-induced phospholipidosis for drug development and safety assessment. Toxicol Appl Pharmacol. 279:467–476.

Lord, S. J., Velle, K. B., Mullins, R. D., and Fritz-Laylin, L. K. 2020. SuperPlots: Communicating reproducibility and variability in cell biology. J Cell Biol. 219:e202001064

Loregger, A., Raaben, M., Nieuwenhuis, J., Tan, J., Jae, L. T., van den Hengel, L. G., Hendrix, S., van den Berg, M., Scheij, S., Song, J. Y., Huijbers, I. J., Kroese, L. J., Ottenhoff, R., van Weeghel, M., van de Sluis, B., Brummelkamp, T., and Zelcer, N. 2020. Haploid genetic screens identify SPRING/C12ORF49 as a determinant of SREBP signaling and cholesterol metabolism. Nat Commun. 11:1128.

Lu, A., Wawro, P., Morgens, D. W., Portela, F., Bassik, M. C., and Pfeffer, S. R. 2018. Genome-wide interrogation of extracellular vesicle biology using barcoded miRNAs. Elife. 7:e41460.

Lu F, Liang Q, Abi-Mosleh L, Das A, De Brabander JK, Goldstein JL, Brown MS. Identification of NPC1 as the target of U18666A, an inhibitor of lysosomal cholesterol export and Ebola infection. Elife. 2015 Dec 8;4:e12177. doi: 10.7554/eLife.12177. PMID: 26646182; PMCID: PMC4718804.

McCauliff LA, Langan A, Li R, et al. 2019. Intracellular cholesterol trafficking is dependent upon NPC2 interaction with lysobisphosphatidic acid. Elife. 2019;8:e50832.

Meneses-Salas, E., García-Melero, A., Kanerva, K., Blanco-Muñoz, P., Morales-Paytuvi, F., Bonjoch, J., Casas, J., Egert, A., Beevi, S. S., Jose, J., Llorente-Cortés, V., Rye, K. A., Heeren, J., Lu, A., Pol, A., Tebar, F., Ikonen, E., Grewal, T., Enrich, C., and Rentero, C. 2020. Annexin A6 modulates TBC1D15/Rab7/ StARD3 axis to control endosomal cholesterol export in NPC1 cells. Cell Mol. Life Sci. 77:2839–2857.

Morgens, D. W., Deans, R. M., Li, A., and Bassik, M.C. 2016. Systematic comparison of CRISPR/Cas9 and RNAi screens for essential genes. Nat Biotechnol. 34:634–636.

Morgens, D. W., Wainberg, M., Boyle, E. A., Ursu, O., Araya, C. L., Tsui, C. K., Haney, M. S., Hess, G. T., Han, K., Jeng, E. E., Li, A., Snyder, M. P., Greenleaf, W. J., Kundaje, A., and Bassik, M. C. 2017. Genome-scale measurement of off-target activity using Cas9 toxicity in high-throughput screens. Nat Commun. 8:15178.

Naito, T., Ercan, B., Krshnan, L., Triebl, A., Koh, D., Wei, F. Y., Tomizawa, K., Torta, F. T., Wenk, M. R., and Saheki, Y. 2019. Movement of accessible plasma membrane cholesterol by the GRAMD1 lipid transfer protein complex. Elife 8:e51401.

Newton, J., Palladino, E., Weigel, C., Maceyka, M., Gräler, M. H., Senkal, C. E., Enriz, R. D., Marvanova, P., Jampilek, J., Lima, S., Milstien, S., and Spiegel, S. 2020. Targeting defective sphingosine kinase 1 in Niemann-Pick type C disease with an activator mitigates cholesterol accumulation. J. Biol.Chem. 295:9121–9133.

Pentchev PG. 2004. Niemann-Pick C research from mouse to gene. Biochim. Biophys. Acta. 1685:3–7.

Pfeffer, S.R. 2019. NPC intracellular cholesterol transporter 1 (NPC1)-mediated cholesterol export from lysosomes. J Biol Chem. 294:1706–1709.

Rosenbaum, A.I., Zhang, G., Warren, J.D., et al. 2010. Endocytosis of beta-cyclodextrins is responsible for cholesterol reduction in Niemann-Pick type C mutant cells. Proc. Natl. Acad. Sci. U S A 107:5477–82.

Saha P, Shumate JL, Caldwell JG, Elghobashi-Meinhardt N, Lu A, Zhang L, Olsson NE, Elias JE, Pfeffer SR. 2020. Inter-domain dynamics drive cholesterol transport by NPC1 and NPC1L1 proteins. Elife. 2020 May 15;9:e57089.

Sakamaki, J. I., Wilkinson, S., Hahn, M., et al. 2017. Bromodomain Protein BRD4 Is a Transcriptional Repressor of Autophagy and Lysosomal Function. Mol Cell 66:517–532.e9. https://doi.org/10.1016/j.molcel.2017.04.027

Salanga, C. M., and Salanga, M. C. 2021. Genotype to Phenotype: CRISPR Gene Editing Reveals Genetic Compensation as a Mechanism for Phenotypic Disjunction of Morphants and Mutants. Int J Mol Sci. 22: 3472.

Sandhu J, Li S, Fairall L, Pfisterer SG, Gurnett JE, Xiao X, Weston TA, Vashi D, Ferrari A, Orozco JL, Hartman CL, Strugatsky D, Lee SD, He C, Hong C, Jiang H, Bentolila LA, Gatta AT, Levine TP, Ferng A, Lee R, Ford DA, Young SG, Ikonen E, Schwabe JWR, Tontonoz P. 2018. Aster Proteins Facilitate Nonvesicular Plasma Membrane to ER Cholesterol Transport in Mammalian Cells. Cell 175(2):514–529.e20.

Scott, C.C., Vossio, S., Vacca, F., Snijder, B., Larios, J., Schaad, O., Guex, N, Kuznetsov, D., Martin, O., Chambon, M, Turcatti, G., Pelkmans, L. and Gruenberg, J. 2015 Wnt directs the endosomal flux of LDL-derived cholesterol and lipid droplet homeostasis. EMBO Rep 16:741–752.

Solomon, L. A., Podder, S., He, J., et al. 2017. Coordination of Myeloid Differentiation with Reduced Cell Cycle Progression by PU.1 Induction of MicroRNAs Targeting Cell Cycle Regulators and Lipid Anabolism. Mol Cell Biol 37:e00013–17.

Strauss, K., Goebel, C., Runz, H., et al. 2010. Exosome secretion ameliorates lysosomal storage of cholesterol in Niemann-Pick type C disease. J Biol Chem 285:26279–26288.

Stuffers, S., Sem Wegner, C., Stenmark, H., et al. 2009. Multivesicular endosome biogenesis in the absence of ESCRTs. Traffic 10:925–937.

Tan, J., Cook, E., van den Berg, M., et al. 2019. Differential use of E2 ubiquitin conjugating enzymes for regulated degradation of the rate-limiting enzymes HMGCR and SQLE in cholesterol biosynthesis. Atherosclerosis 281:137–142.

Thelen, A. M., and Zoncu, R. 2017. Emerging Roles for the Lysosome in Lipid Metabolism. Trends Cell Biol. 27:833–850.

Trajkovic, K., Hsu, C., Chiantia, S., et al. 2008. Ceramide triggers budding of exosome vesicles into multivesicular endosomes. Science 319:1244–1247.

Trinh MN, Brown MS, Goldstein JL, et al. 2020. Last step in the path of LDL cholesterol from lysosome to plasma membrane to ER is governed by phosphatidylserine. Proc. Natl. Acad. Sci. U S A 117:18521–18529.

Ugrankar R, Bowerman J, Hariri H, et al. 2019. Drosophila Snazarus Regulates a Lipid Droplet Population at Plasma Membrane-Droplet Contacts in Adipocytes. Dev. Cell 50:557–572.e5.

van den Boomen, D.J.H., Sienkiewicz, A., Berlin, I. et al. 2020. A trimeric Rab7 GEF controls NPC1-dependent lysosomal cholesterol export. Nat. Commun. 11, 5559.

Wang, B., and Tontonoz, P. 2018. Liver X receptors in lipid signalling and membrane homeostasis. Nat Rev Endocrinol. 14:452–463.

Yamanaka, T., Tosaki, A., Kurosawa, M., et al., 2016. Genome-wide analyses in neuronal cells reveal that upstream transcription factors regulate lysosomal gene expression. FEBS J. 283:1077–1087.

Youn, D. Y., Xiaoli, A. M., Pessin, J. E., et al. 2016. Regulation of metabolism by the Mediator complex. Biophys Rep. 2:69–77.

